# ACTIVATION OF THE ARYL HYDROCARBON RECEPTOR REDUCES VISCERAL HYPERSENSITIVITY IN A MOUSE MODEL OF COLITIS IN REMISSION

**DOI:** 10.64898/2026.11.04.763733

**Authors:** Yasaman B. Habibyan, Colin Mackenzie, Armaan Mohan, Cristiane H. Baggio, Noah Gordon, Md Abu Naser Mohon, Timothy E. Shutt, Manon Defaye, Christina S. Thornton, Braedon McDonald, Christophe Altier, Keith A. Sharkey, Yasmin Nasser

**Affiliations:** Department of Medicine, Snyder Institute for Chronic Diseases, Cumming School of Medicine, University of Calgary, Calgary, AB, Canada; Department of Critical Care Medicine, Snyder Insitute for Chronic Diseases, Cumming School of Medicine, University of Calgary, Calgary, AB, Canada; Departments of Biochemistry and Molecular Biology and Medical Genetics, Hotchkiss Brain Institute, Snyder Institute for Chronic Diseases and Alberta Children’s Hospital Research Institute, Cumming School of Medicine, Calgary, AB, Canada; Department of Microbiology, Immunology and Infectious Diseases, Snyder Institute for Chronic Diseases, Cumming School of Medicine, University of Calgary, Calgary, AB, Canada; Department of Physiology and Pharmacology, Snyder Institute for Chronic Diseases and Hotchkiss Brain Institute, Cumming School of Medicine, Calgary, AB, Canada

**Keywords:** visceral pain, indole-3-pyruvic acid, aryl hydrocarbon receptor, Inflammatory Bowel Disease, DSS colitis

## Abstract

The aryl hydrocarbon receptor (AhR) is expressed on gut-projecting transient receptor potential vanilloid-1 (TRPV1)^+^ nociceptors, but whether it has a role in regulating abdominal pain remains to be determined. Approximately 50% of inflammatory bowel disease (IBD) patients experience chronic abdominal pain, even when their inflammatory disease is in remission. Previous work has shown that tryptophan metabolism to indole derivatives is reduced in IBD patients. Since microbial tryptophan metabolites (indole derivatives) serve as ligands for AhR, we hypothesized that reduced AhR activation contributes to enhanced pain sensitivity. We used a post-colitis mouse model of visceral hypersensitivity to examine the role of AhR as a mediator of nociception. AhR activation using both natural and synthetic agonists reduced visceral hypersensitivity in both male and female post-colitis mice. Conversely, pharmacological inhibition of AhR blocked the antinociceptive effects of indoles. Mechanistically, AhR activation by indole-3-pyruvic acid (IPyA) increased the transcription of the inward rectifying potassium channel Kir2.2, in TRPV1^+^ neurons and markedly reduced neuronal activation within the dorsal horn of the spinal cord. Metagenomic analysis revealed sex differences in indole-producing bacteria, with control males exhibiting enriched abundance of *Akkermansia muciniphila* and *Alistipes* compared to post-colitis males and female controls. These findings demonstrate that oral IPyA supplementation in post-colitis mice attenuates visceral hypersensitivity in part through activation of nociceptor AhR, providing insights that support the therapeutic targeting of AhR to treat abdominal pain in IBD patients in remission.

## Introduction

Inflammatory bowel diseases (IBD) are chronic inflammatory disorders of the gastrointestinal (GI) tract. IBD disease activity is characterized by relapsing periods of remission and flares (1). Remission refers to a lack of active inflammation determined by endoscopic evaluation; flares refer to periods of active disease and inflammation, with associated symptoms such as acute abdominal pain, rectal bleeding and diarrhea. The primary objective of medical therapy in IBD is to induce and maintain remission of inflammatory disease, as there is no cure. However, approximately 50% of IBD patients experience chronic abdominal pain, even when their inflammatory disease is in remission (2, 3). Unfortunately, there are few treatments tailored toward chronic abdominal pain in the absence of active inflammation in IBD, due in part to a gap in knowledge regarding underlying pathophysiological mechanisms. Consequently, up to 24% of IBD patients experiencing chronic pain receive no treatment (4).

Recently, the gut microbiota has emerged as a critical modulator of visceral hypersensitivity, a condition where pain-sensing nerves, known as nociceptors, become sensitized. Microbial products have been shown to both directly stimulate nociceptors, through corresponding receptors (5–7), or stimulate them indirectly via immune or endocrine mechanisms (8, 9). We have previously demonstrated that visceral hypersensitivity can be directly transferred from mice that had recovered from acute colitis (post-colitis mice) via a fecal microbiota transplant (FMT) to naïve germ-free mice (10). Several clinical studies have reported altered microbial profiles in patients suffering from chronic and/or recurrent visceral pain, including patients with IBD (11–16). Notably, indoles—the primary byproducts of bacterial tryptophan (Trp) metabolism—are significantly decreased in IBD patients during both flares and remission (17, 18). These data suggest that a change in the microbial milieu in IBD patients in remission or mice that have recovered from colitis contributes to chronic visceral pain.

The aryl hydrocarbon receptor (AhR) is a candidate indole receptor that can bind to numerous indole derivatives with varying affinity (19). AhR was originally identified as a xenobiotic receptor, however, recent research has revealed that AhR signalling is involved in intestinal homeostasis and epithelial barrier integrity (20). Furthermore, AhR has been shown to be expressed throughout the peripheral and central nervous system (21, 22). A 2019 study utilizing single-cell RNA sequencing mapped seven functional classes of visceral sensory dorsal root ganglia (DRG) neurons projecting to the mouse colon, and demonstrated that AhR is highly expressed in five out of these seven classes, including transient receptor potential vanilloid-1 (TRPV1) expressing nociceptors (21).

Here we used a post-colitis mouse model of visceral hypersensitivity (10, 23) to examine the role of AhR as a mediator of nociception. AhR activation via both natural and synthetic agonists decreased visceral hypersensitivity in both male and female post-colitis mice. Conversely, pharmacological inhibition of AhR blocked the antinociceptive effects of indoles. Mechanistically, AhR activation by indole-3-pyruvic acid (IPyA) increased the transcription of the inward rectifying potassium channel Kir2.2, in TRPV1^+^ neurons and markedly reduced neuronal activation within the dorsal horn of the spinal cord. Together, these findings demonstrate that oral IPyA supplementation in post-colitis mice attenuates visceral hypersensitivity in part by activating nociceptor AhR.

## Methods

### Sex as a biological variable

All experiments were performed in age-matched male and female mice to evaluate any potential sex-specific differences.

### Animals

5-week-old male and female C57BL/6 mice were obtained from Jackson Laboratories (Bar Harbor, ME, USA) and allowed to rest for 1 week before experiments. Mice were cohoused in groups of 3 in ventilated cages (Green Line GM500, SealSafe Plus, Techniplast) with unrestricted access to standard chow and water. Cages were housed in controlled environments on a 12-hour light and 12-hour dark cycle. Mice within a single cage were randomly assigned to the various experimental groups to account for confounding variables due to cohousing. All mouse studies were conducted after approval from the University of Calgary Health Sciences Animal Care Committee (animal use protocol AC24-0134) and were conducted in accordance with the guidelines established by the Canadian Council on Animal Care

### Dextran Sulphate Sodium (DSS) induced colitis and disease evaluation

Male and female mice were given either 2.5% (wt./vol) dextran sulphate sodium (DSS; 40,000 MW, ThermoFisher, Waltham, MA, USA) in drinking water or drinking water alone for 5 days. Following DSS treatment, all mice were given regular drinking water and allowed to recover for 5 weeks. This is referred to as the post-colitis DSS mouse model of visceral hypersensitivity (23). Bodyweight changes and disease activity index were recorded daily until recovery (day 13) and every 3 days thereafter until day 42.

Weight loss, stool consistency, and bleeding were monitored daily to evaluate the severity of DSS-induced colitis. The Disease Activity Index (DAI) score was calculated using a previously published 12-point scale system (24). The following formula was used: DAI = (weight loss score) + (stool consistency score) + (hematochezia score).

**Table 1.** Scoring system for the DSS Disease Activity Index (DAI)

| Score | Weight loss (%) | Stool consistency | Hematochezia |
|---|---|---|---|
| 0 | 0 | Normal | Absence |
| 1 | 1-5 |  |  |
| 2 | 6-10 | Loose stool |  |
| 3 | 11-15 |  |  |
| 4 | >15 | Diarrhea | Presence |

Adapted from Park et al. (24).

Colon length was assessed at the time of death. The entire mouse colon was collected, and the total length was measured. A 1 cm section of the distal colon was collected and fixed overnight in 4% paraformaldehyde (PFA). Tissues were embedded and sectioned by the Core Pathology Lab at the University of Calgary Libin Cardiovascular Unit. For microscopic damage scoring, tissue slides were stained with hematoxylin and eosin for histologic analysis. The microscopic damage score was calculated using a 6-point score (25).

### Estrus Cycle Synchronization

To control for the effects of sex hormones on visceral sensitivity, female mice were synchronized using vasectomized males. One week before visceral sensitivity measurements, three cycling female mice were cohoused with a single vasectomized male. Visceral sensitivity measurements were then performed in the diestrus or metestrus stage. On the day of visceral sensitivity measurements, the presence of the diestrus or metestrus stage was confirmed by analyzing the leukocyte count compared to cornified and nucleated cells (10).

### β-naphthoflavone (BNF), Indole-3-pyruvic acid (IPyA), and GNF351 administration

BNF (Sigma-Aldrich, Oakville, ON, CA, Cat. No. N36633-1G) was prepared following our previously published methods (26). The BNF solution was administered intraperitoneally (IP) at 40 mg/kg for 3 days before visceral sensitivity measurement. Control mice received a vehicle (IP) injection of corn oil. Indole-3-pyruvic acid (IPyA; Sigma-Aldrich, Oakville, Cat. No. I7017-5G) was mixed into water at a concentration of 20mg/mL immediately prior to gavage. Mice were gavaged once daily with IPyA (200 mg/kg) for 3 days prior to visceral sensitivity evaluation. Control mice received a gavage of water only (10 ml/kg). GNF351 stock solution (Sigma-Aldrich, Cat. No. 182707) was diluted in DMSO (20 mg/ml) and stored at-20°C. GNF351 stock solution was diluted into fresh saline (0.1 mg/ml) on each day of injection, and 10 μg was administered IP. Control mice received a vehicle (IP) injection of 0.05% DMSO in saline. For all pharmacological studies mice within a single cage were randomly assigned to either vehicle or drug groups

### Visceral Motor Reflex to Colorectal Distention

Visceral sensitivity was assessed using the visceral motor response (VMR) to colorectal distention. Mice were anesthetized and two electrodes were implanted into the right abdominal external oblique muscles and exteriorized through the back of the neck. To prevent the mice from removing the electrodes, a plastic tube was sutured to the skin at the exteriorization site. The mice were then allowed to recover for 48 hours prior to visceral sensitivity measurement.

On the day of testing, mice were individually restrained in 50 mL falcon tubes. A 10.5 mm balloon catheter was placed 0.5 cm deep into the rectum of each mouse. Mice were allowed to rest for thirty minutes before starting measurements. The rectal balloon was inflated at 15, 30, 45, and 60 mmHg pressures, each for 10 seconds; mice were allowed to rest for 5 minutes between each distention. Electrodes were connected to a Bio Amplifier which signals an electromyogram acquisition system to record and display the response (ADInstruments, Colorado Springs, CO, USA). The electromyographic activity was recorded from the abdominal contractions, and the visceromotor response was calculated using LabChart 7 software (ADInstruments).

### RNA isolation and PCR

The distal colon and the visceral innervating DRGs (T10-L1; L6-S1) were isolated and stored in RNAlater (Invitrogen, Burlington, ON, CA, Cat. No. AM7021) at 4°C until RNA isolation. RNA was isolated using the RNeasy Mini Kit (Qiagen, Toronto, ON, CA, Cat. No. 74104). RNA quantity and quality were measured using NanoDrop 2000c spectrophotometer (ThermoFisher). Purified RNA was reverse transcribed into cDNA using a high-capacity reverse transcription kit (Applied Biosystems, Waltham, MA, USA). The cycle conditions for RT were as follows: 25°C for 10 minutes, 37°C for 120 minutes, and 85°C for 5 seconds.

Samples were amplified in a 20 µL reaction mix using Applied Biosystems SYBR® Green MasterMix. *Cyp1a1* expression was measured using the primer sequence Forward: CATCACAGACAGCCTCATTGAGC and Reverse: CTCCACGAGATAGCAGTTGTGAC. b-Actin (*Bact*) expression was measured using the primer sequence Forward: CATTGCTGACAGGATGCAGAAGG and Reverse: TGCTGGAAGGTGGACAGTGAGG. Gene expression was normalized to the housekeeping gene (*Bact*) and the 2^-ΔΔCT^ method was used to measure relative expression.

### Long-read metagenomic sequencing, functional metagenomics and data analysis

Two to three pellets of stool were collected from male and female control and post-colitis mice at day 42 and stored at-80°C until DNA isolation. Bacterial DNA was isolated from samples using the Allprep PowerFecal Pro DNA/RNA kit (Qiagen). A Qubit fluorometer (ThermoFisher) was used to quantify the concentration of DNA from all samples.

Metagenomic gDNA samples were barcoded using the Oxford Nanopore Technologies Ligation sequencing gDNA – Native Barcoding Kit 96 V14 (SQK-NBD114.96) following the protocol version NBE_9171_v114_revQ_23Dec2024 according to the manufacturer’s direction with no modifications. The barcoded gDNA library was sequenced using the Oxford Nanopore Technologies PromethION 2 Solo system on a FLO-PRO114M flow cell for 120 hours. Basecalling was completed within MinKNOW (v.25.05.14) using Dorado (v.7.9.8) with the Super-accurate basecalling 400bps model, with barcodes trimmed and a minimum Q-score of 10 (Oxford Nanopore Technologies). Taxonomy was assigned from the FASTQ files containing reads above the Q-score threshold using the Epi2Me (Oxford Nanopore Technologies) app (v.5.2.5) with the pipeline wf-metagenomics running Kraken2 (v.2.1.3) with default parameters using the prebuilt Kraken2 database PlusPF-8. The Kraken2 report output files were converted into a biom format using the tool kraken-biom (v.1.2.0). The biom file was imported into R as a phyloseq object using the phyloseq (v.1.54.2) package. Relative abundance plots were computed and visualized using the microviz (v.0.13.0) package. β-diversity was calculated using Bray-Curtis distances and PCoA ordination was plotted using the microviz package. α-diversity was calculated using the microbiome (v.1.32.0) package and subsequently plotted using ggplot2 (v.4.0.3) with the Wilcoxon rank-sums generated using the ggpubr (v.0.6.3) package. Linear discriminant analysis effect size (LEFSe) was calculated using the microbiomeMarker (v.1.13.2) package.

### Immunofluorescence and Confocal microscopy – spinal cord

Mice were perfused with phosphate buffered saline (PBS) followed by 4% PFA to wash out blood and fix tissues. Isolated lumbosacral (L6-S1) spinal cord sections were fixed in 4% PFA overnight at 4°C. The tissues were then washed in PBS and placed to dehydrate in 30% sucrose at 4°C. Tissues were embedded using optimal cutting temperature (OCT) solution (VWR International, Cat. No. 95057-838). The samples were then sectioned (16 μm), with 5 sections per slide on Superfrost Plus slides (ThermoFisher).

To assess neuronal activation in the dorsal horn of the spinal cord, tissues were first blocked for 1 hour in goat serum in 0.3% Triton x-100 solution. Samples were then incubated overnight at 4°C with primary antibody mix in 5% fetal bovine serum (FBS) (c-Fos, 1:500, Abcam, Cat. No. ab190289, Waltham, MA, USA; NeuN, 1:1000, Sigma-Aldrich, Cat. No. MAB377). The next day, sections were washed 3x in PBS and then incubated at room temperature with secondary antibody in 5% fetal bovine serum (FBS) (goat anti-rabbit Alexa 488, 1:1000, Invitrogen, Cat. No. A11034; goat anti-mouse Alexa 568, 1:1000, Invitrogen, Cat. No. A11031). Images were visualized and taken with a IXplore IX83 SpinSR (Evident Scientific, Waltham, MA, USA). For cFos counts, at least 3 sections per mouse were imaged, and the right and left dorsal horns were counted and then averaged. Means between experimental groups were compared using ordinary one-way analysis of variance (ANOVA) with Tukey post hoc test.

### RNAscope In Situ Hybridization – dorsal root ganglion neurons (DRGs)

Mice were perfused with PBS followed by 4% PFA to wash out blood and fix tissues. Isolated lumbosacral (L6-S1) DRGs were fixed in 4% PFA overnight at 4°C. The tissues were then washed in PBS and placed to dehydrate in 30% sucrose at 4°C. Tissues were embedded in OCT solution. The samples were then sectioned (10 μm), with 8 sections per slide on Superfrost Plus slides (ThermoFisher) and stored at-80°C. The RNAscope assay was performed over two days. On the first day of the assay slides were first washed in 1x PBS to remove OCT, and then baked for 60 minutes at 60°C. The sections were then postfixed in prechilled 4% PFA at 4°C for 15 minutes. The sections were next dehydrated by 5-minute washes in 50% ethanol, 70% ethanol, and 100% ethanol x2. Barriers were drawn around sections with a hydrophobic pen (Cat. No. H-4000, Vector Labs, Newark, CA, USA) and sections were incubated with hydrogen peroxide (Cat. No. 322335, ACDBio, Newark, CA, USA) for 10 minutes at room temperature (RT). Sections were rinsed twice with distilled water prior to incubation with protease III for 30 minutes at 40°C in the ACD HybEZTM Oven (ACDBio). The RNAscope assay was then performed according to ACDBio’s instructions. Slides were then mounted using ProLong Gold antifade reagent (Invitrogen, Cat. No. P36934). The probes used were Mm-Trpv1 (ACD, Cat. No. 313331), Mm-Ahr (ACD, Cat. No. 452091-C2), Mm-Cyp1a1 (ACD, Cat. No. 464661-C3), and Mm-Kcnj12 (ACD, Cat. No. 525171-C3) and were labelled using TSA Vivid Fluorophores 520 (ACD, Cat. No. 323271), 570 (ACD, Cat. No. 323272) and 650 (ACD, Cat. No. 323273). Images were visualized and taken with a FLUOVIEW FV4000 Confocal Laser Scanning Microscope (Evident Scientific). RNAscope quantification was performed using QuPath software, following the recommendations provided by the ACD manufacturer. The expected dot size was set to was approximately 0.3 µm^2^, with acceptable sizes ranging from 0.1 µm^2^ to 0.7 µm^2^. Means between experimental groups were compared using two-way ANOVA with Tukey’s multiple comparison test.

## Statistical Analysis

Data analysis was performed using GraphPad Prism 10.0 software (GraphPad, La Jolla, CA, USA) with the exception of long-read sequencing, which was analyzed as outlined above. All data presented is expressed as the mean ± standard error mean. The Shapiro-Wilk test of normality was first used to evaluate the distribution of the data. If the data followed a normal distribution, the mean was compared using parametric tests; otherwise, nonparametric methods were used to analyze the data. If a normal, or Gaussian distribution was confirmed, an ordinary one-way/two-way ANOVA with Tukey post hoc test was applied for multiple groups with one or two variables, respectively. Alternatively, the Kruskal-Wallis nonparametric test with Dunn’s post hoc test was used. A p value<0.05 was considered significant.

## Results

### Activation of AhR reduces visceral hypersensitivity in post-colitis mice

To examine the role of AhR signalling in post-inflammatory chronic visceral hypersensitivity, we used the established DSS post-colitis mouse model (10, 23). In this model, male and female mice are given 2.5% DSS for 5 days and allowed to recover for 5 weeks (Figure 1A). We first tested whether oral supplementation of a microbial Trp metabolite could attenuate post-inflammatory visceral hypersensitivity. Previous studies in mouse models have used oral administration of various microbial Trp metabolites, to study their downstream effects *in vivo* (29–31). Aoki *et al.* demonstrated that oral supplementation of IPyA activated AhR in the colon to alleviate inflammation in a mouse model of acute DSS colitis (29). Thus, for these studies, we used the natural AhR agonist IPyA given orally. (Figure 1A).

**Figure 1.**
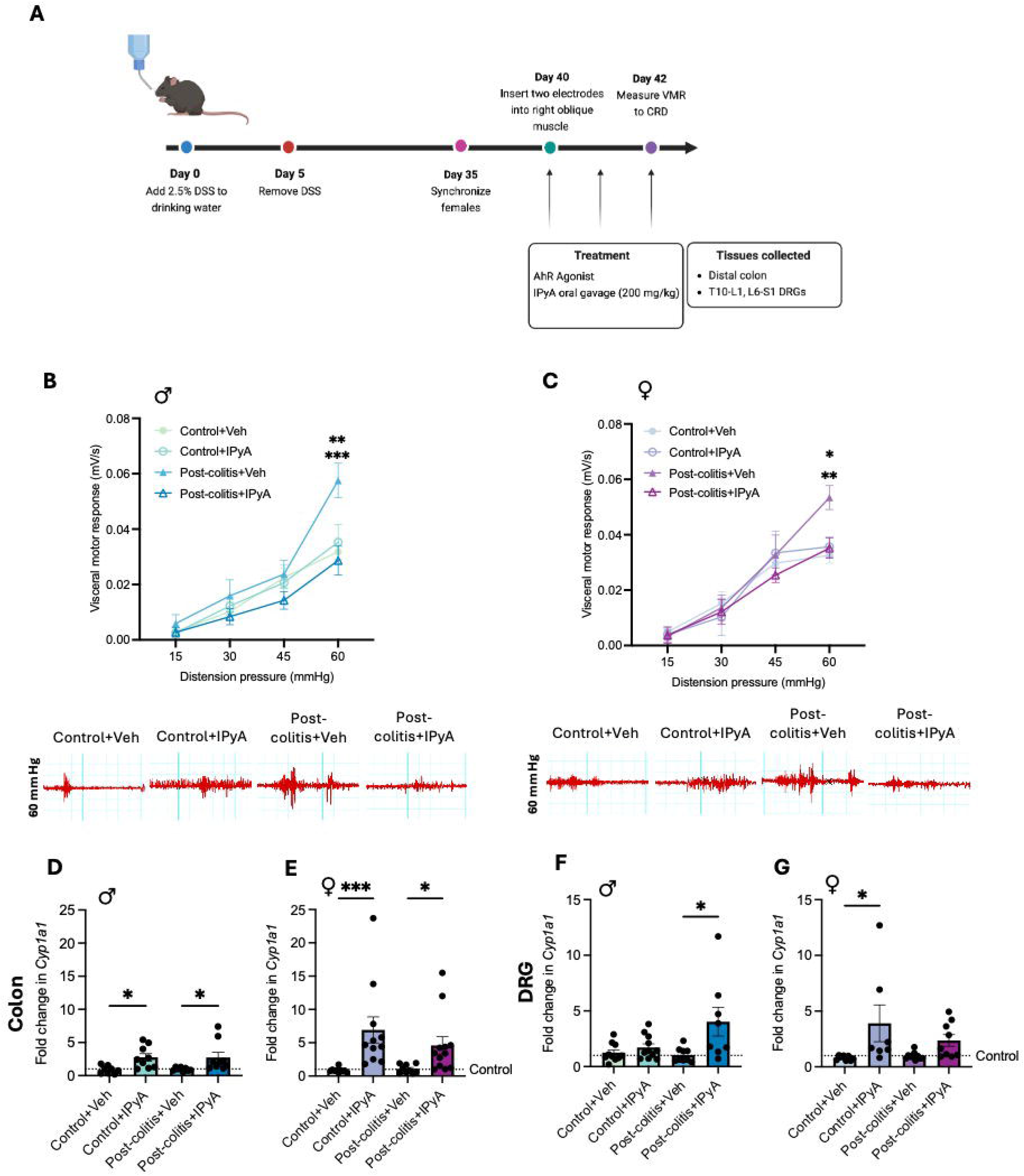
Treatment with IPyA decrease visceral hypersensitivity in post-colitis male and female mice. (A) Male and female mice were given 2.5% DSS or water for 5 days and allowed to recover for 5 weeks. At day 35, a vasectomized male was introduced to synchronize females. Electrodes were implanted at day 40 and visceral pain assessment was performed at day 42 followed by euthanasia of animals. From day 40-42 mice were administered either AhR agonist, IPyA or vehicle control. Following euthanization, the distal colon and T10-L1, L6-S1 DRGs were collected. (B) Visceral sensitivity was assessed using the visceromotor response to colorectal distention in control and post-colitis males treated with IPyA or vehicle. 60 mmHg: Control+Veh vs Post-colitis+Veh p<0.001; Control+IPyA vs Post-colitis+Veh p<0.01; Post-colitis+Veh vs Post-colitis+IPyA p<0.0001. Two-way ANOVA with Tukey’s multiple comparison test. Data expressed as mean ± SEM. n=7-10 mice per group. (C) Visceral sensitivity was assessed using the visceromotor response to colorectal distention in control and post-colitis females treated with IPyA or vehicle. 60 mmHg: Control+Veh vs Post-colitis+Veh p<0.01; Control+IPyA vs Post-colitis+Veh p<0.05; Post-colitis+Veh vs Post-colitis+IPyA p<0.05. Data expressed as mean ± SEM. n=4-6 mice per group. Distal colon tissue from males (E) and females (F) treated with IPyA was collected to measure the relative expression of *Cyp1a1*. DRGs were collected from the T10-L1 and L6-S1 segments from males (G) and females (H) treated with IPyA was collected to measure the relative expression of *Cyp1a1*. *p<0.05; **p<0.01; ***p<0.001; ****p<0.0001. Relative mRNA expression compared to the housekeeping gene b-actin was calculated to produce ΔCT values. The control ΔCT values were averaged, and each ΔCT from all samples was compared to this value to generate the ΔΔCT. 2^-ΔΔCT^ was calculated and plotted to show fold change expression. Data expressed as mean ± SEM. n=7-12 per group.

Both male and female mice treated with DSS displayed weight loss and increased disease activity that resolved following the cessation of DSS (Supplemental Figure 1). Colon length and histological staining were used to confirm the resolution of inflammation at the time of behaviour testing (Supplemental Figure 1). IPyA (200 mg/kg) was administered by oral gavage for 3 days leading up to behaviour testing, with the last dose given 1 hour prior to testing (Figure 1A).

In all experiments, visceral sensitivity measurements were performed in both males and diestrus/metestrus females, as visceral sensitivity is known to vary across the estrus cycle. The diestrus and metestrus stages of the estrus cycle are considered low-estrogen phases of the estrus cycle (32). Visceral sensitivity was examined using the VMR to colorectal distention (Figure 1A). Both male and female post-colitis mice exhibited increased visceral sensitivity when compared to their sex-matched control counterparts (Figure 1B, 1C). Oral supplementation with IPyA decreased visceral sensitivity back to baseline levels in both male and female post-colitis mice (Figure 1B, 1C). IPyA administration to control animals did not reduce visceral sensitivity (Figure 1B, 1C).

To confirm successful activation of AhR in our model, we used RT-qPCR to examine mRNA levels of, a downstream target of AhR commonly used to determine AhR activation (33). A segment of the distal colon and visceral innervating DRGs (T10-L1; L6-S1) were collected to confirm *Cyp1a1* upregulation. Administration of IPyA upregulated *Cyp1a1* mRNA levels in the distal colon of both control and post-colitis male and female mice (Figure 1D, 1E). In the DRGs of male mice, IPyA administration led to upregulation of *Cyp1a1* mRNA in only post-colitis but not control groups (Figure 1F). In females, IPyA administration resulted in *Cyp1a1* upregulation only in the control state (Figure 1G).

Despite seeing a significant reduction in visceral hypersensitivity, oral supplementation of IPyA did not induce significant upregulation of *Cyp1a1* transcripts in the DRGs of post-colitis females (or control males); we wondered whether stimulation of AhR with a potent synthetic agonist would yield similar results. To examine this, we used a synthetic AhR agonist, beta-naphthoflavone (BNF; 40 mg/kg), which was administered IP for 3 days leading up to behavioural testing, with the last dose given 1 hour prior to testing (Figure 2A). As before, male and female mice given DSS displayed weight loss and increased disease activity that resolved following the cessation of DSS (Supplemental Figure 2). Colon length and histological staining were used to confirm the resolution of inflammation at the time of behaviour testing (Supplemental Figure 2).

**Figure 2.**
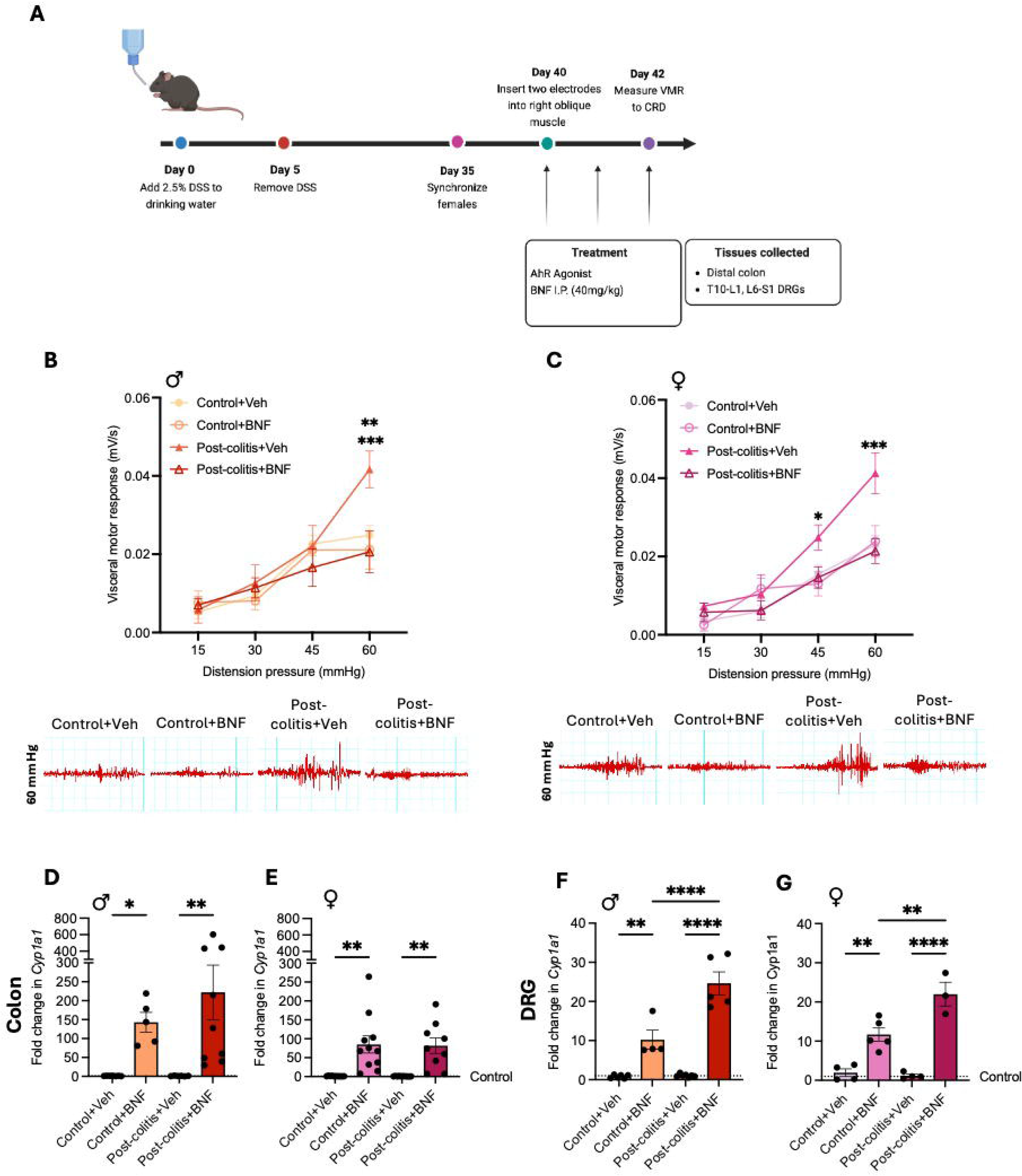
Treatment with BNF decreases visceral hypersensitivity in post-colitis male and female mice. (A) Male and female mice were given 2.5% DSS or water for 5 days and allowed to recover for 5 weeks. At day 35, a vasectomized male was introduced to synchronize females. Electrodes were implanted at day 40 and visceral pain assessment was performed at day 42 followed by euthanasia of animals. From day 40-42 mice were administered either AhR agonist, BNF or vehicle control. Following euthanization, the distal colon and T10-L1, L6-S1 DRGs were collected. (B) Visceral sensitivity was assessed using the visceromotor response to colorectal distention in control and post-colitis males treated with BNF or vehicle. 60 mmHg: Control+Veh vs Post-colitis+Veh p<0.01; Control+BNF vs Post-colitis+Veh p<0.01; Post-colitis+Veh vs Post-colitis+BNF p<0.01. Two-way ANOVA with Tukey’s multiple comparison test. Data expressed as mean ± SEM. n=5-7 mice per group. (C) Visceral sensitivity was assessed using the visceromotor response to colorectal distention in control and post-colitis females treated with BNF or vehicle. 45 mmHg: Control+BNF vs Post-colitis+Veh p<0.05. 60 mmHg: Control+Veh vs Post-colitis+Veh p<0.001; Control+BNF vs Post-colitis+Veh p<0.001; Post-colitis+Veh vs Post-colitis+BNF p<0.001. Two-way ANOVA with Tukey’s multiple comparison test. Data expressed as mean ± SEM. n=6-7 mice per group. Distal colon tissue from males (D) and females (D) treated with BNF was collected to measure the relative expression of *Cyp1a1*. DRGs were collected from the T10-L1 and L6-S1 segments from males (F) and females (G) treated with BNF was collected to measure the relative expression of *Cyp1a1*. *p<0.05; **p<0.01; ***p<0.001; ****p<0.0001. Relative mRNA expression compared to the housekeeping gene b-actin was calculated to produce ΔCT values. The control ΔCT values were averaged, and each ΔCT from all samples was compared to this value to generate the ΔΔCT. 2^-ΔΔCT^ was calculated and plotted to show fold change expression. Data expressed as mean ± SEM. n=3-11 per group.

As seen with IPyA, treatment with BNF reduced post-inflammatory visceral hypersensitivity back to control levels in both sexes (Figure 2B, 2C). Moreover, administration of BNF in control animals did not further reduce visceral sensitivity (Figure 2B, 2C). Administration of BNF strongly upregulated *Cyp1a1* mRNA levels in the distal colon of both control and post-colitis males and females (Figure 2D, 2E). Moreover, we observed that BNF treatment significantly upregulated *Cyp1a1* transcripts in the DRGs of control and post-colitis males and females (Figure 2F, 2G). Interestingly, in the DRGs, relative *Cyp1a1* upregulation was significantly higher in the post-colitis groups of both sexes (Figure 2F, 2G). This may indicate that AhR activity is reduced in the post-colitis groups, allowing for a greater relative increase when stimulated. Ultimately, we observed comparable effectiveness between IPyA and BNF in reducing post-inflammatory visceral hypersensitivity in our model (Figures 1B, 1C, 2B, 2C).

### Pharmacological inhibition of AhR blocks the antinociceptive effect of IPyA

In addition to AhR, microbial Trp metabolites have been shown to act on the pregnane X receptor (PXR) (34). Thus, to confirm that IPyA acts through only AhR to exert the antinociceptive effect we observed, we pretreated mice with the AhR antagonist, GNF351 (10 μg) 30 minutes prior to oral gavage of IPyA (200 mg/kg) for 3 days leading up to behaviour testing, with the last combined dose given 1 hour prior to testing (Supplemental Figure 3). Both male and female mice given DSS in their drinking water presented with weight loss and increased disease activity that resolved following the termination of DSS (Supplemental Figure 4 and 5). Colon length and histological staining were used to confirm the resolution of inflammation at the time of behaviour testing of all animals (Supplemental Figure 4 and 5).

Both male and female post-colitis mice exhibited increased visceral sensitivity when compared to their sex-matched control counterparts (Figure 3). Treatment with IPyA in male and female post-colitis mice attenuated visceral hypersensitivity (Figure 3C, 3G). Notably, inhibition of AhR by GNF351 in post-colitis male and female mice blocked the antinociceptive effect of IPyA (Figure 3C, 3G). However, inhibition of AhR in post-colitis mice did not further increase visceral hypersensitivity (Figure 3C, 3G). Interestingly, in control male and female mice, inhibition of AhR with GNF351 alone resulted in visceral hypersensitivity (Figure 3A, 3E). Importantly, IPyA gavage did not reduce visceral hypersensitivity in control male and female mice treated with GNF351, suggesting that IPyA acts through AhR to decrease visceral hypersensitivity (Figure 3).

**Figure 3.**
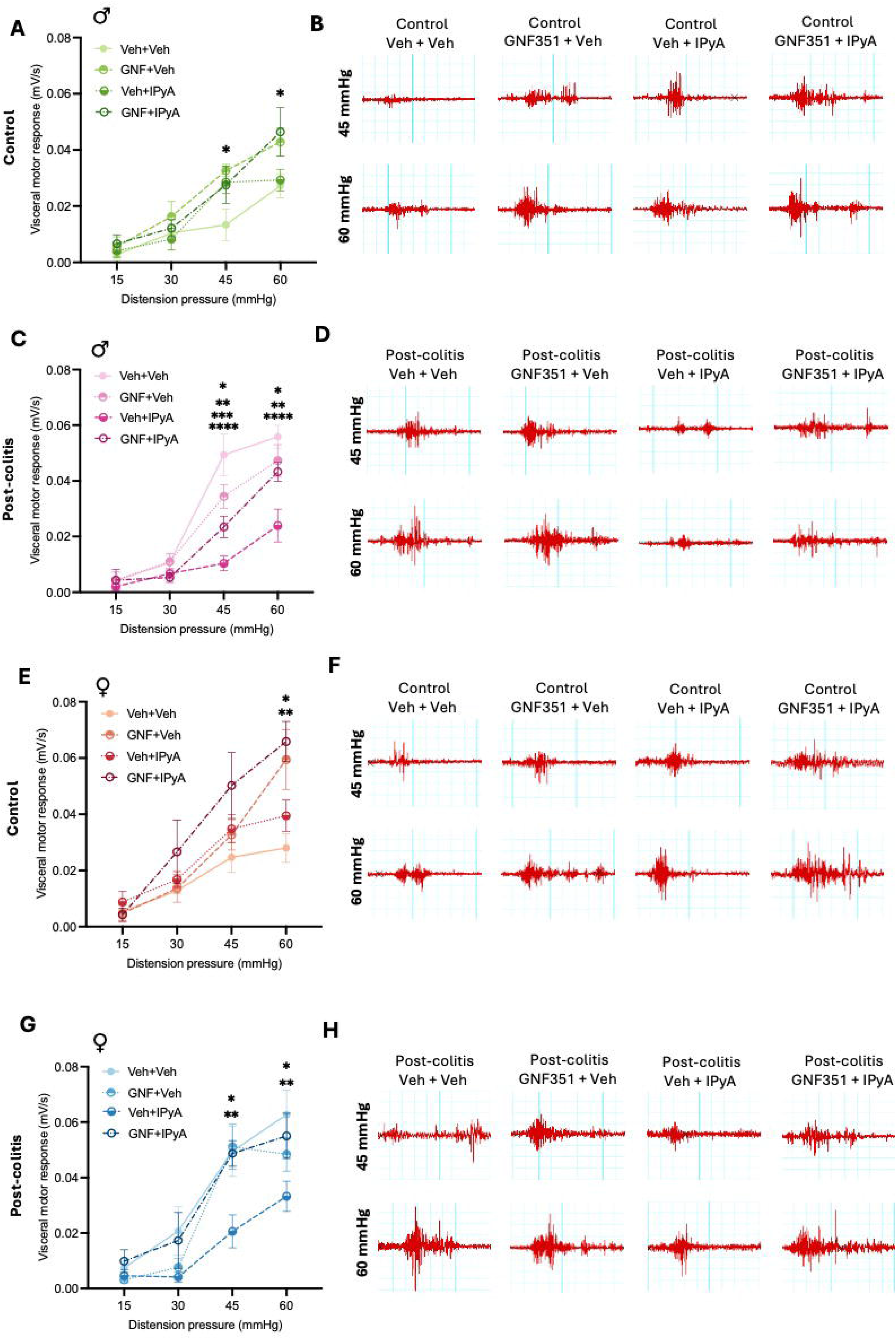
Pharmacological inhibition of AhR blocks the effect of IPyA in post-colitis males and females and increases visceral sensitivity at baseline. (A) Visceral sensitivity was assessed using the visceromotor response to colorectal distention in control males treated with GNF351 and IPyA. Two-way ANOVA with Tukey’s multiple comparison test. Data expressed as mean ± SEM. n=4-5 per group – see Supplemental Tables 1 and 2 for significance. (B) Representative traces at 45 mmHg and 60 mmHg for all control male groups. (C) Visceral sensitivity was assessed using the visceromotor response to colorectal distention in post-colitis males treated with GNF351 and IPyA. Two-way ANOVA with Tukey’s multiple comparison test. Data expressed as mean ± SEM. n=5-6 per group – see Supplemental Tables 1 and 2 for significance. (D) Representative traces at 45 mmHg and 60 mmHg for all post-colitis male groups. (E) Visceral sensitivity was assessed using the visceromotor response to colorectal distention in control females treated with GNF351 and IPyA. Two-way ANOVA with Tukey’s multiple comparison test. Data expressed as mean ± SEM. n=4-5 per group – see Supplemental Tables 3 and 4 for significance. (F) Representative traces at 45 mmHg and 60 mmHg for all control female groups. (G) Visceral sensitivity was assessed using the visceromotor response to colorectal distention in post-colitis females treated with GNF351 and IPyA. Two-way ANOVA with Tukey’s multiple comparison test. Data expressed as mean ± SEM. n=6 per group – see Supplemental Tables 3 and 4 for significance. (H) Representative traces at 45 mmHg and 60 mmHg for all post-colitis female groups.

### IPyA reduces neuronal activation in the spinal cord of post-colitis animals

Following colorectal distention, cFos positive neurons were quantified in the lumbosacral (L6-S1) spinal cord of animals. Post-colitis male and female mice had increased neuronal activation in the spinal cord when compared to control groups, which was restored to control levels with IPyA (Figure 4). Cotreatment with GNF351 and IPyA in post-colitis animals resulted in similar neuronal activation as the post-colitis vehicle groups, indicating that IPyA acts through AhR in our model (Figure 4). Supplementation of IPyA to control male and female mice did not reduce neuronal activation below baseline levels.

**Figure 4.**
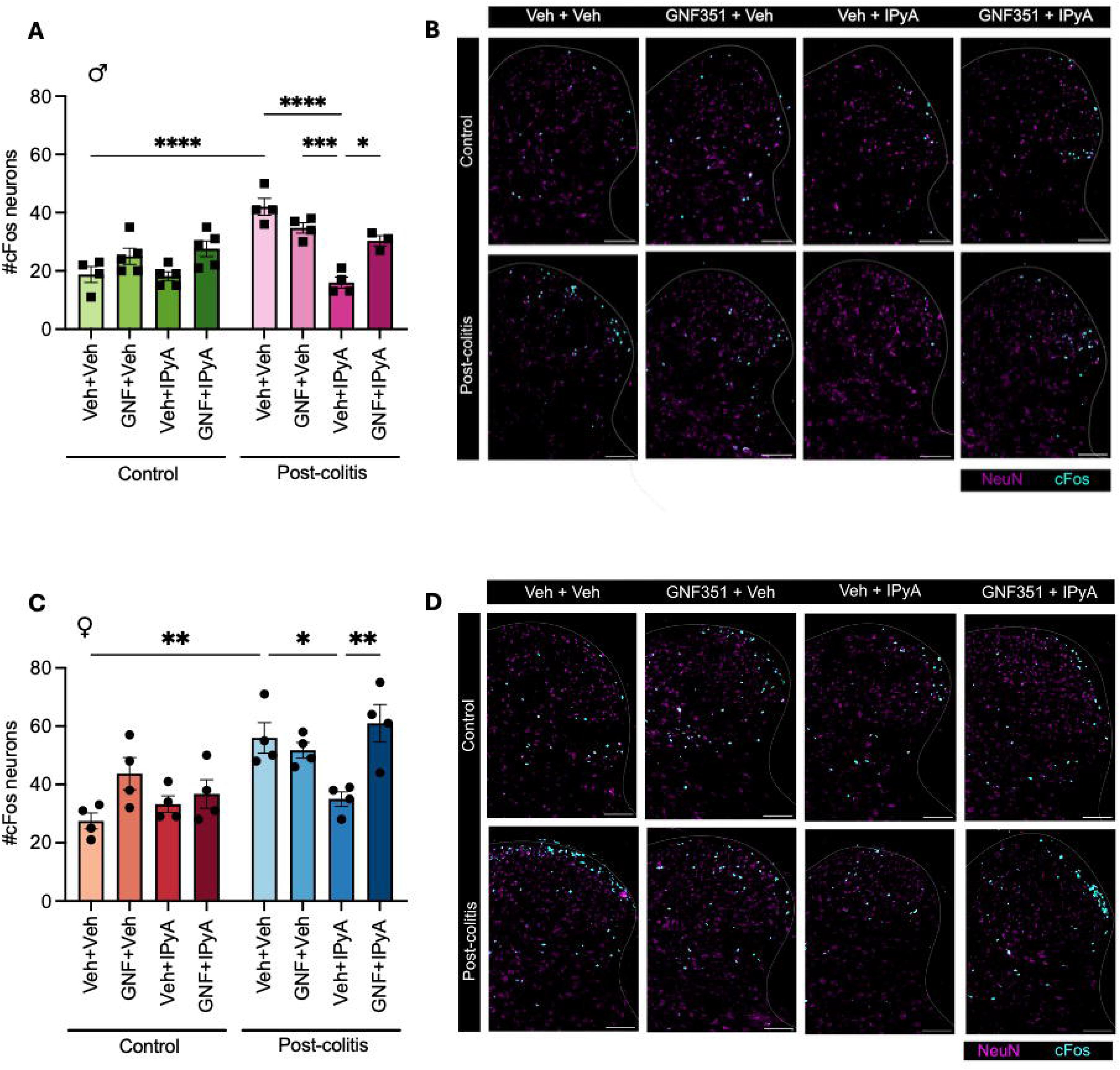
IPyA supplementation decreases neuronal activation in the lumbosacral spinal cord of post-colitis males and females. (A) Neuronal activation was measured in the dorsal horn of the lumbosacral (L6/S1) spinal cord of males. Activated neurons were counted where NeuN and cFos signal colocalized. Data expressed as mean ± SEM. n=3-4 per group. *p<0.05; ***p<0.001; ****p<0.0001. (B) Representative images of cFos (cyan) and NeuN (magenta) in the dorsal horn of the spinal cord of male animals. Scale bar: 200 µm. (C) Neuronal activation was measured in the dorsal horn of the lumbosacral (L6/S1) spinal cord of females. Activated neurons were counted where NeuN and cFos signal colocalized. Data expressed as mean ± SEM. n=3-4 per group. *p<0.05; **p<0.01. (D) Representative images of cFos (cyan) and NeuN (magenta) in the dorsal horn of the spinal cord of female animals. Scale bar: 200 µm.

**Figure 5.**
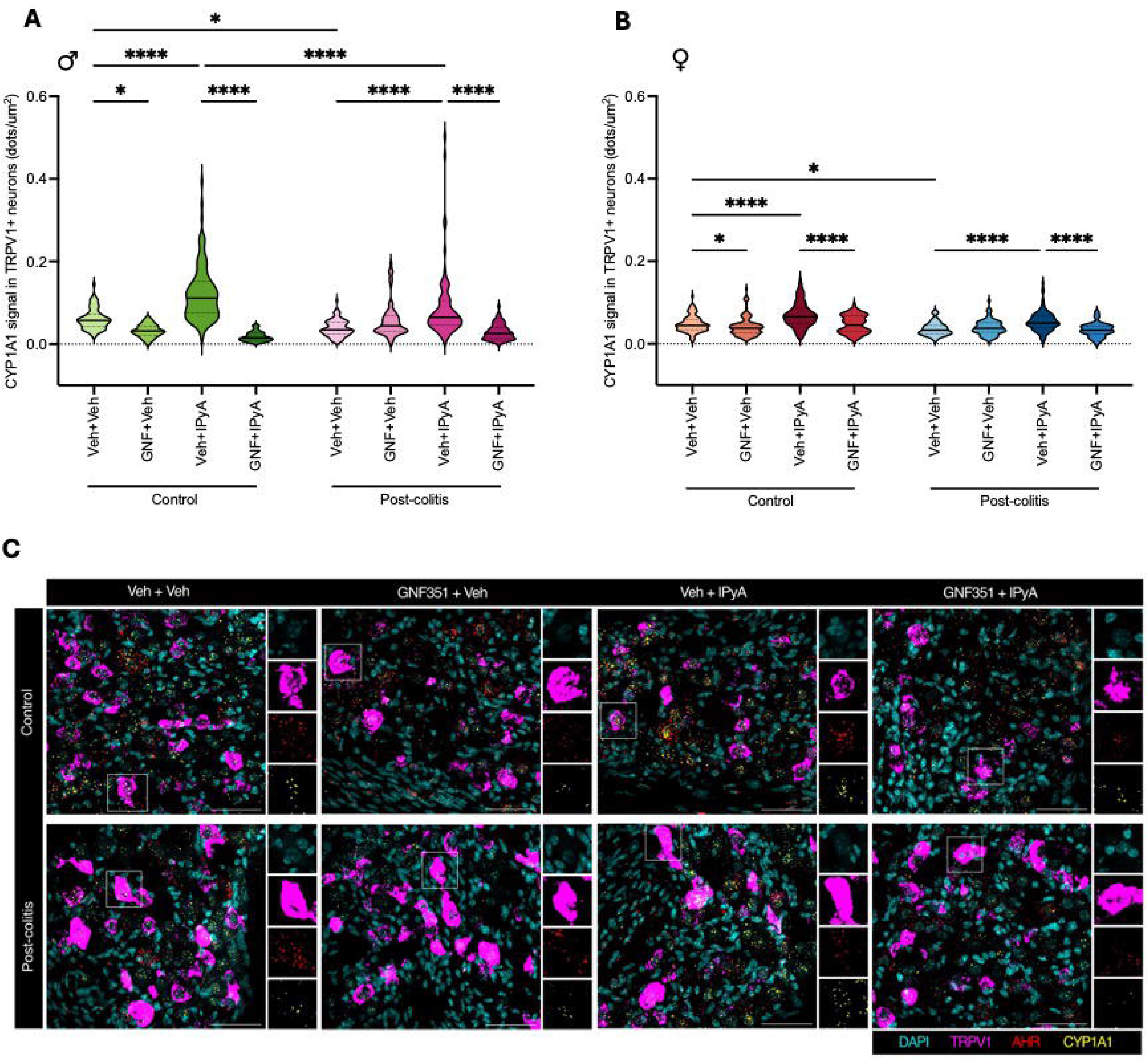
IPyA supplementation increases CYP1A1 transcripts in TRPV1^+^ DRG neurons of males and females. (A-B) *Cyp1a1* transcripts were quantified in TRPV1^+^ DRG neurons of male and female mice. *Cyp1a1* signal was determined by calculating the number of estimated dots/areas. A dot was considered positive if the size was approximately 0.3µm^2^, with acceptable sizes ranging from 0.1 µm^2^ to 0.7 µm^2^. Two-way ANOVA with Tukey’s multiple comparison test. Data expressed as mean ± SEM. n=3-4 mice per group. *p<0.05; **p<0.01; ***p<0.001; ****p<0.0001. (C) Representative RNAscope images from males showing expression of *Trpv1* transcripts (magenta), *Ahr* transcripts (red) and *Cyp1a1* transcripts (yellow) with DAPI staining (cyan) in lumbosacral (L6/S1) DRGs. Scale bar: 50 µm.

### Treatment with IPyA increased AhR activation in TRPV1^+^ DRG neurons

Because the DRG comprises a heterogeneous cell population, we reasoned that whole-tissue qRT-PCR might fail to detect localized transcriptional changes in *Cyp1a1* within TRPV1⁺ DRG neurons. To selectively quantify AhR activity in TRPV1^+^ DRG neurons, *Cyp1a1* transcripts were quantified using RNAscope, a fluorescent in situ hybridization (FISH) assay. A decrease in *Cyp1a1* transcripts was observed in TRPV1^+^ DRG neurons of post-colitis males and females, when compared to sex-matched control groups, indicating a decrease in AhR activity within TRPV1^+^ nociceptors in these mice (Figure 5A-C). Inhibition of AhR by GNF351 did not further decrease *Cyp1a1* transcripts in post-colitis groups (Figure 5A-C). Treatment with IPyA in both male and female post-colitis mice led to the restoration of *Cyp1a1* transcripts to control levels (Figure 5A-C). However, pretreatment with GNF351 prior to IPyA blocked this effect in post-colitis male and female mice (Figure 5A-C). In control male and female mice, inhibition of AhR, using GNF351 led to a decrease in *Cyp1a1* transcripts, suggesting AhR is active within TRPV1^+^ nociceptors at baseline. Administration of IPyA to both male and female control mice further increased *Cyp1a1* transcripts in TRPV1^+^ nociceptors. However, pretreatment with GNF351 prior to IPyA blocked this effect (Figure 5A-C).

### Treatment with IPyA increased Kcnj12 expression in TRPV1^+^ DRG neurons of post-colitis males and females

Recent work has shown that *Ahr* expression in enteric neurons is positively correlated with *Kcnj12* expression, which encodes Kir2.2, an inward rectifying potassium channel (35). Increased expression of Kir2.2 has been shown to decrease neuronal excitability (36). As we observed decreased neuronal activation in the dorsal horn of post-colitis male and female mice treated with IPyA, we examined *Kcnj12* transcription in TRPV1^+^ nociceptors in the lumbosacral (L6-S1) DRGs. In both post-colitis males and females, there was a decrease in *Kcnj12* transcripts in TRPV1^+^ DRG neurons when compared to sex-matched control groups (Figure 6A-C). Treatment with IPyA in post-colitis males and females increased *Kcnj12* expression relative to sex-matched control and post-colitis vehicle levels. However, when post-colitis males and females were pretreated with GNF351 prior to IPyA administration, *Kcnj12* signal was comparable to that of sex-matched post-colitis vehicle groups. Treatment with IPyA in control females, but not males, further increased *Kcnj12* transcripts compared to control vehicle groups. Moreover, in control females, pretreatment with GNF351 prior to IPyA administration resulted in decreased *Kcnj12* transcripts compared to control IPyA females; this was not seen in males. Finally, in control males treated with GNF351, *Kcnj12* signal was decreased compared to control vehicle males, however this was not seen in females (Figure 6A-C).

**Figure 6.**
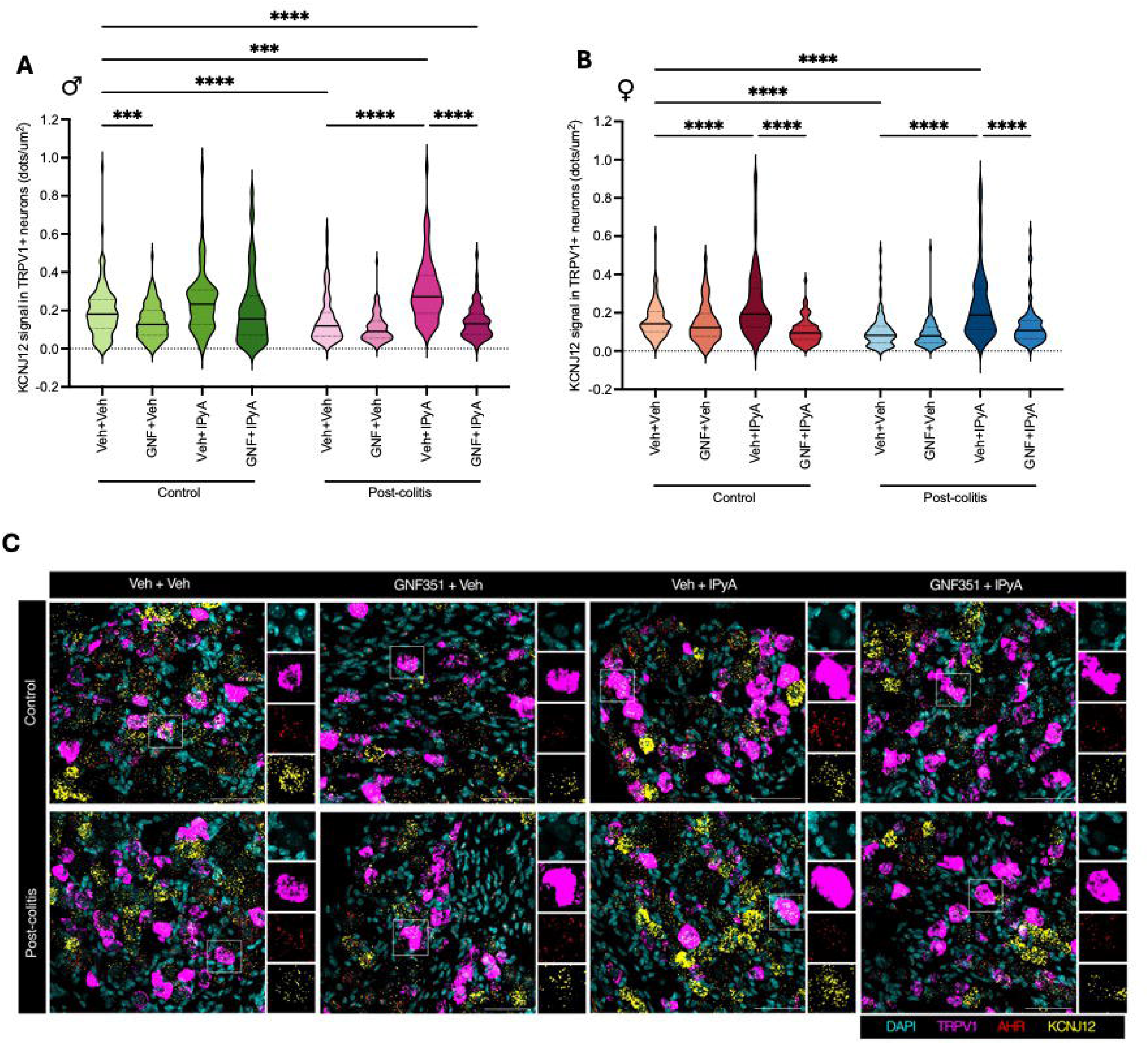
IPyA supplementation increases KCNJ12 transcripts in TRPV1^+^ DRG neurons of males and females. (A-B) *Kcnj12* transcripts were quantified in TRPV1+ DRG neurons of male and female mice. *Kcnj12* signal was determined by calculating the number of estimated dots/areas. A dot was considered positive if the size was approximately 0.3µm^2^, with acceptable sizes ranging from 0.1 µm^2^ to 0.7 µm^2^. Two-way ANOVA with Tukey’s multiple comparison test. Data expressed as mean ± SEM. n=3-4 mice per group. *p<0.05; **p<0.01; ****p<0.0001. (C) Representative RNAscope images from males showing expression of *Trpv1* transcripts (magenta), *Ahr* transcripts (red) and *Kcnj12* transcripts (yellow) with DAPI staining (cyan) in lumbosacral (L6/S1) DRGs. Scale bar: 50 µm.

### Microbial communities involved in tryptophan metabolism are decreased in the post-colitis model of visceral hypersensitivity in a sex-specific manner

As we could reverse visceral hypersensitivity in the post-colitis state with AhR activation, we wondered whether the gut microbiota of post-colitis males and females exhibited a loss in indole-producing bacterial species. We observed that the composition of male but not female post-colitis and control microbiota were significantly altered (Figure 7A, 7B, 7C). No differences in the alpha diversity of control and post-colitis groups were observed in either sex (Supplemental Figure 6). To better visualize changes between control and post-colitis groups we conducted differential abundance analysis. We observed that compared to post-colitis males, control males had an enrichment of *Akkermansia muciniphila* and several members of the genus *Alistipes* (Figure 7D). Moreover, we observed that compared to control females, control males had an enrichment of *A. muciniphila* and several members of the genus *Alistipes* (Figure 7E). Relative abundance plots were generated to visualize the alteration of *A. muciniphila* abundance in control versus post-colitis males and females (Supplemental Figure 7). Notably, when compared to males, the relative abundance of *A. muciniphila* was reduced and more variable within female cohorts (Supplemental Figure 7).

**Figure 7.**
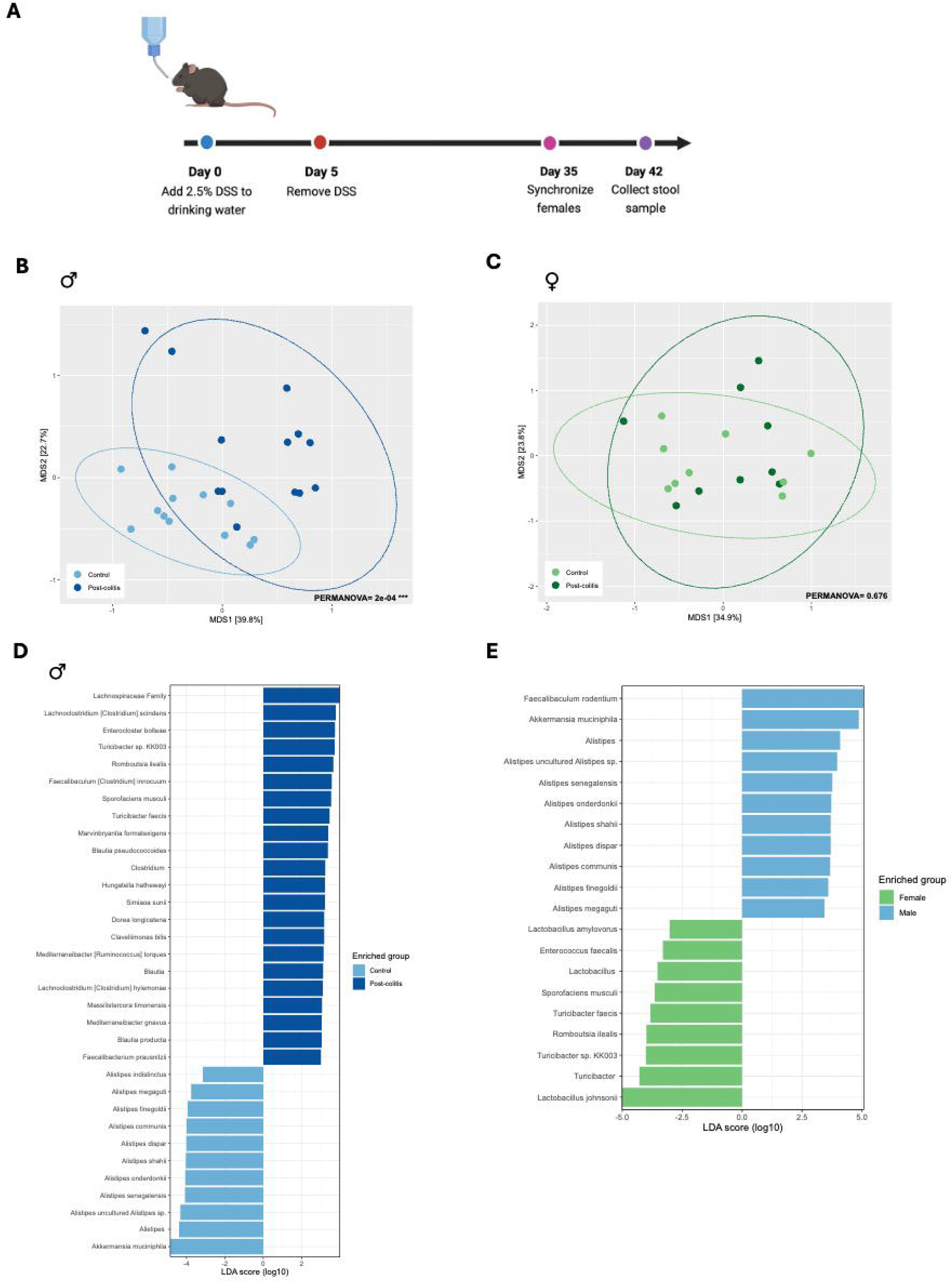
The fecal microbiota of control and post-colitis males but not females is significantly altered. (A) Male and female mice were given 2.5% DSS or water for 5 days and allowed to recover for 5 weeks. At day 35, a vasectomized male was introduced to synchronize females. Stool was collected from mice at day 42. Beta diversity was measured by generating a Bray-Curtis dissimilarity matrix (MDS) for males (B) and females (C). PERMANOVA statistical analysis was used to compare groups, n=9-13 mice per group. Linear discriminant analysis (LDA) effect size (LEFSe) was calculated to identify differences between control and post-colitis males (D) and males and females (E). Bacteria that reached a LDA score (log10) > 2.0 were selected as significant, n=9-13 mice per group.

## Discussion

Approximately 50% of patients with IBD will experience chronic abdominal pain at some point following diagnosis (37–39). Despite this, the mechanisms which drive this visceral sensitization are not well understood, leading to a lack of therapeutic targets to manage symptoms. In recent years, the gut microbiota has emerged as a key regulator in the gut-brain axis (5–7, 9). Here, we demonstrate that oral supplementation with IPyA—a microbial tryptophan metabolite— activates AhR within TRPV1^+^ nociceptors, suppressing neuronal excitability and central activation to alleviate visceral hypersensitivity in post-colitis mice of both sexes.

Patients with IBD display persistent alterations to the composition and functionality of their gut microbiota (40). Importantly, these alterations endure irrespective of disease activity (41). Altered microbial Trp metabolism has been implicated in both animal models and patients with IBD (18, 27, 42). Both murine and human colitis are associated with the selective depletion of indole and indole-derived metabolites (17). Additionally, patients with IBD in remission have reduced fecal indole levels, with a reduced capacity to activate AhR *in vitro* (18).

We characterized the fecal microbiota in a post-colitis model of visceral hypersensitivity, revealing distinct, sex-dependent alterations. Control males specifically displayed a marked enrichment of *A. muciniphila* and *Alistipes* compared to post-colitis males—a shift absent in females—consistent with baseline male enrichment of these taxa. Recent findings have shown that oral supplementation of *A. muciniphila* attenuates colonic inflammation by enhancing microbial Trp metabolism and activating AhR signalling (28). Similarly, stratification of patients with IBD by AhR activity established an elevated abundance of *Alistipes* in individuals with high AhR activity (27). Together, these findings suggest that the post-colitis depletion of bacterial species that make indoles, such as *A. muciniphila* and *Alistipes*, in males, but not females, disrupts downstream Trp pathways to drive persistent male visceral hypersensitivity. While potential microbiota transfer during cohousing with vasectomized males may have mitigated differences in female groups—representing a limitation of our study design—further examination of potential sex differences in microbial Trp metabolism in IBD remains a valuable avenue for future investigation.

AhR signalling has been implicated in the pathophysiology of active IBD. Various studies have shown that activation of AhR using either microbes or metabolites, such as IPyA, can regulate intestinal function and attenuate intestinal inflammation (28, 29, 43). Recently, Rondeau *et al.* showed that a high Trp diet in specific-pathogen free mice increased microbial Trp metabolites, leading to increased AhR activation in the colonic epithelium, which provided protection against DSS colitis (27). Due to the anti-inflammatory and restorative properties of IPyA and AhR, we administered IPyA during the last 3 days of our model, after active colonic inflammation had fully resolved. This timing was intentionally chosen to decouple the resolution of colitis from visceral hypersensitivity mitigation, allowing us to isolate and evaluate the specific role of post-inflammatory AhR signaling in the visceral pain pathway.

To our knowledge, we are the first to examine the role of AhR signalling as a target for mediating post-inflammatory visceral hypersensitivity. We examined whether IPyA, a microbially produced AhR activating ligand, could ameliorate post-colitis visceral hypersensitivity. Oral IPyA supplementation effectively reduced visceral hypersensitivity in post-colitis males and females, highlighting a potentially safe, microbiome-targeted therapeutic strategy. Using a potent synthetic AhR agonist, BNF, we demonstrated that activation of AhR was also sufficient to reduce post-inflammatory visceral hypersensitivity in males and females. However, despite the successful use of synthetic AhR agonists in mouse models, they do not serve as viable therapeutic candidates, due to concerns regarding their immunotoxicity and carcinogenicity (44, 45).

Using RNAscope, we demonstrated that oral supplementation of IPyA stimulated AhR signalling in TRPV1^+^ DRG neurons, through increased *Cyp1a1* transcripts. We confirmed the specificity of this pathway using the AhR antagonist GNF351, which has previously been shown to inhibit IPyA-induced activation of AhR *in vivo* (29). Consistently, we found that pharmacological inhibition of AhR successfully blocked IPyA-induced *Cyp1a1* upregulation. Collectively, these findings establish that oral IPyA ameliorates post-inflammatory visceral hypersensitivity in both sexes through targeted AhR activation in TRPV1^+^ nociceptors, which can be blocked by pharmacological AhR inhibition.

While our study focused specifically on nociceptor AhR activation in pain transmission, we also assessed broad AhR activation in the distal colon via qRT-PCR, observing that IPyA upregulated *Cyp1a1* transcripts in both sexes. Because AhR is widely expressed throughout the intestinal epithelium and immune compartments (46, 47), these local cells may also contribute to the visceral nociceptive pathway. Future studies should investigate the specific colonic cell types activated by oral IPyA supplementation in the post-colitis model of visceral hypersensitivity.

Peripheral sensitization—characterized by a lowered threshold and heightened responsiveness of primary afferent nerve endings—is driven by inflammatory mediators binding to receptors that modulate ion channels such as TRPV1, ultimately enhancing nerve fibre excitability and causing hyperalgesia (48–50). To investigate the downstream mechanism by which AhR signaling modulates nociception, we assessed neuronal activation within the lumbosacral spinal cord following colorectal distention. Lumbosacral nociceptors traveling via sacral pelvic nerves primarily innervate the distal colon and rectum (51). Distention triggered robust dorsal horn activation in post-colitis mice, consistent with previous work (52). Specifically, oral IPyA supplementation suppressed lumbosacral dorsal horn activation in post-colitis mice, which, was blocked by pharmacological AhR inhibition. Together, these data demonstrate that microbial-targeted AhR activation alleviates visceral hypersensitivity by dampening central propagation of peripheral pain signals.

Previous work has shown that AhR activity in enteric neurons positively correlated with the transcription of *Kcnj12* (35). *Kcnj12* encodes Kir2.2, an inward rectifying potassium channel (35). Kir2 channels are essential modulators of neuronal excitability as they maintain the resting membrane potential of neurons (36). In our study, we showed that *Kcnj12* transcription was reduced in the TRPV1^+^ DRG neurons of post-colitis male and female mice. This observed decrease in *Kcnj12* transcription provides mechanistic insight into post-inflammatory visceral hypersensitivity. Excitingly, activation of AhR by IPyA in post-colitis males and females upregulated the transcription of *Kcnj12* in TRPV1^+^ nociceptors to baseline levels. While these findings suggest that AhR signaling attenuates visceral hypersensitivity by reducing neuronal excitability, direct electrophysiological validation remains an important future direction.

Pharmacological inhibition of AhR in healthy control mice of both sexes was sufficient to induce visceral hypersensitivity, indicating that baseline AhR signaling plays a role in regulating the visceral pain pathway physiologically. Interestingly, GNF351 treatment in control males reduced *Kcnj12* transcripts within TRPV1^+^ nociceptors, which was not observed in control female groups. Conversley, IPyA supplementation significantly upregulated AhR activation in the visceral innervating DRGs of baseline control females, but not males. Correspondingly, metagenomic analysis revealed that control males had an enrichment of bacteria that produce indoles, including *A. muciniphila* and *Alistipes*, relative to control females. Previous work by Bayrer *et al.* investigating the role of enterochromaffin (EC) cells and isovalerate, a microbially produced short-chain fatty acid in visceral hypersensitivity, revealed sex-differences exist under physiological conditions. They established that the EC cell-mucosal afferent signalling pathway was active at baseline in females, but not males (9). Together, these findings demonstrate that at baseline, sex biased differences in visceral pain due to microbial signalling may exist through host-microbiome interactions.

Although the role of AhR signalling in nociception is not well characterized, published work has demonstrated that AhR signalling is involved in the microbiota-gut-brain axis. AhR is broadly expressed throughout the central and peripheral nervous system, including the cerebral cortex, astrocytes, glial cells, and dorsal root ganglia (21, 22). Importantly, mice with a global AhR knockout develop progressive neuropathic pain, whereas AhR activation exerts neuroprotective and anti-inflammatory effects (22). Furthermore, AhR signaling has been linked to behavioral changes: the AhR/IL-22 pathway alleviates anxiety-like behaviors in post-infectious IBS models (53), and microbial indole derivatives mitigate mood disorders in an AhR-dependent manner (54). Together with our findings, these studies establish AhR as a compelling therapeutic target for disorders of the microbiota-gut-brain axis.

In summary, our findings demonstrate that targeting microbial–AhR signalling offers a novel therapeutic strategy for mitigating post-inflammatory visceral hypersensitivity. Activation of the microbial-AhR pathway via microbial metabolite supplementation potentially serves as a safe and cost-effective alternative to current therapies available to IBD patients in remission with chronic abdominal pain. We show that oral supplementation of the microbial metabolite, IPyA activates AhR in TRPV1^+^ primary afferent neurons. AhR activation is accompanied by reduced neuronal activation in the lumbosacral spinal cord and upregulation of *Kcnj12* transcription in nociceptors, resulting in reduced neuronal excitability in both male and female post-colitis mice. Although baseline differences in the microbial communities involved in tryptophan metabolism in males and females show sex-specific differences, IPyA-mediated AhR activation effectively reverses visceral hypersensitivity across both sexes. Taken together, these results highlight the microbial–AhR pathway as a promising target for managing chronic post-inflammatory abdominal pain in patients with IBD in remission.

## Supplemental Figure Legends

**Supplemental Figure 1.** (A) Weight changes in control and post-colitis male mice. (B) Disease activity index in post-colitis male mice. (C) Colonic length changes in control and post-colitis male mice treated with IPyA. (D) Representative photomicrographs of colonic cross sections of males treated with IPyA. No microscopic evidence of inflammation was seen. Scale bar: 100 µm. (E) Weight changes in control and post-colitis female mice. (F) Disease activity index in post-colitis female mice. (G) Colonic length changes in control and post-colitis female mice treated with IPyA. (H) Representative photomicrographs of colonic cross sections of females treated with IPyA. No microscopic evidence of inflammation was seen. Scale bar: 100 µm.

**Supplemental Figure 2.** (A) Weight changes in control and post-colitis male mice. (B) Disease activity index in post-colitis male mice. (C) Colonic length changes in control and post-colitis male mice treated with BNF. (D) Representative photomicrographs of colonic cross sections of males treated with BNF. No microscopic evidence of inflammation was seen. Scale bar: 100 µm. (E) Weight changes in control and post-colitis female mice. (F) Disease activity index in post-colitis female mice. (G) Colonic length changes in control and post-colitis female mice treated with BNF. (H) Representative photomicrographs of colonic cross sections of females treated with BNF. No microscopic evidence of inflammation was seen. Scale bar: 100 µm.

**Supplemental Figure 3.** (A) Male and female mice were given 2.5% DSS or water for 5 days and allowed to recover for 5 weeks. At day 35, a vasectomized male was introduced to synchronize females. Electrodes were implanted at day 40 and visceral pain assessment was performed at day 42 followed by euthanasia of animals. From day 40-42 mice were administered a cotreatment of GNF351, an AhR antagonist with IPyA. Appropriate groups were given vehicle controls. (B) Following euthanasia, the lumbosacral spinal cord and DRGs were collected for immunofluorescence and FISH, respectively. A section of the distal colon was collected for histology.

**Supplemental Figure 4.** (A) Weight changes in control and post-colitis male mice. (B) Disease activity index in post-colitis male mice. (C) Colonic length changes in control and post-colitis male mice cotreated with GNF351 and IPyA. (D) Representative photomicrographs of colonic cross sections of males cotreated with GNF351 and IPyA. No microscopic evidence of inflammation was seen. Scale bar: 100 µm.

**Supplemental Figure 5.** (A) Weight changes in control and post-colitis female mice. (B) Disease activity index in post-colitis female mice. (C) Colonic length changes in control and post-colitis female mice cotreated with GNF351 and IPyA. (D) Representative photomicrographs of colonic cross sections of females cotreated with GNF351 and IPyA. No microscopic evidence of inflammation was seen. Scale bar: 100 µm.

**Supplemental Figure 6.** Alpha Diversity was measured using the Shannon Diversity Index in control and post-colitis males (A) and females (B) and no differences were observed.

**Supplemental Figure 7.** Relative abundance of fecal microbiota from male (A) and female (B) control and post-colitis mice.

## Author Contributions

YBH, CT, BM, KAS, and YN conceptiualized and designed the study; YBH, CM, AM, CB, NG, MANM, and MD performed the experiments; YBH, CM analyzed and interpreted the data; YN acquired funding. TES, CR, BM, KAS, and YN provided supervision. YBH drafted the manuscript; YBH, KAS, and YN critically revised the manuscript for important intellectual content. All authors had access to the study data and have reviewed and approved the final manuscript.

## Acknowledgements

The authors acknowledge the Bioinstrumentation Labs and bioimaging access and support from the Live Cell Imaging Laboratory at the Snyder Institute for Chronic Diseases, Cumming School of Medicine, University of Calgary. The authors also acknowledge the Core Pathology Lab in the Libin Cardiovascular institute, Cumming School of Medicine, University of Calgary. Figures were created with BioRender (biorender.com). Portions of this manuscript have been presented in abstract form at Canadian Digestive Diseases Week (CDDW) and Neurogastro 2025.

## Funding

Supported by the Canadian Institutes of Health Research (CIHR - YN), Crohn’s and Colitis Canada/Pfizer Canada Women in IBD Award (YN), CIHR postdoctoral fellowship (MD), the Department of Medicine, Cumming School of Medicine, University of Calgary (YBH), Alberta Graduate Excellence (AGES) Scholarship (YBH), Canadian Association of Gastroenterology/TRIANGLE Canada award (YBH).

## Conflicts of interest

The authors disclose no conflicts of interest.

## Notes

### Competing Interest Statement

The authors have declared no competing interest.

## References

1. Baumgart DC, Carding SR. Inflammatory bowel disease: cause and immunobiology. Lancet. 2007;369(9573):1627–40.

2. Drewes AM, Olesen AE, Farmer AD, Szigethy E, Rebours V, Olesen SS. Gastrointestinal pain. Nat Rev Dis Primers. 2020;6(1):1.

3. Srinath AI, Walter C, Newara MC, Szigethy EM. Pain management in patients with inflammatory bowel disease: insights for the clinician. Therap Adv Gastroenterol. 2012;5(5):339–57.

4. Zeitz J, Ak M, Muller-Mottet S, Scharl S, Biedermann L, Fournier N, et al. Pain in IBD Patients: Very Frequent and Frequently Insufficiently Taken into Account. PLoS One. 2016;11(6):e0156666.

5. Sessenwein JL, Baker CC, Pradhananga S, Maitland ME, Petrof EO, Allen-Vercoe E, et al. Protease-Mediated Suppression of DRG Neuron Excitability by Commensal Bacteria. J Neurosci. 2017;37(48):11758–68.

6. Baker CC, Sessenwein JL, Wood HM, Yu Y, Tsang Q, Alward TA, et al. Protease-Induced Excitation of Dorsal Root Ganglion Neurons in Response to Acute Perturbation of the Gut Microbiota Is Associated With Visceral and Somatic Hypersensitivity. Cell Mol Gastroenterol Hepatol. 2024;18(4):101334.

7. De Palma G, Lynch MD, Lu J, Dang VT, Deng Y, Jury J, et al. Transplantation of fecal microbiota from patients with irritable bowel syndrome alters gut function and behavior in recipient mice. Sci Transl Med. 2017;9(379).

8. Hughes PA, Harrington AM, Castro J, Liebregts T, Adam B, Grasby DJ, et al. Sensory neuro-immune interactions differ between irritable bowel syndrome subtypes. Gut. 2013;62(10):1456–65.

9. Bayrer JR, Castro J, Venkataraman A, Touhara KK, Rossen ND, Morrie RD, et al. Gut enterochromaffin cells drive visceral pain and anxiety. Nature. 2023;616(7955):137–42.

10. Arzamendi MJ, Habibyan YB, Defaye M, Shute A, Baggio CH, Chan R, et al. Sex-specific post-inflammatory dysbiosis mediates chronic visceral pain in colitis. Gut Microbes. 2024;16(1):2409207.

11. Frank MG, Baratta MV, Sprunger DB, Watkins LR, Maier SF. Microglia serve as a neuroimmune substrate for stress-induced potentiation of CNS pro-inflammatory cytokine responses. Brain Behav Immun. 2007;21(1):47–59.

12. Manichanh C, Rigottier-Gois L, Bonnaud E, Gloux K, Pelletier E, Frangeul L, et al. Reduced diversity of faecal microbiota in Crohn’s disease revealed by a metagenomic approach. Gut. 2006;55(2):205–11.

13. Conte MP, Schippa S, Zamboni I, Penta M, Chiarini F, Seganti L, et al. Gut-associated bacterial microbiota in paediatric patients with inflammatory bowel disease. Gut. 2006;55(12):1760–7.

14. Carroll IM, Ringel-Kulka T, Siddle JP, Ringel Y. Alterations in composition and diversity of the intestinal microbiota in patients with diarrhea-predominant irritable bowel syndrome. Neurogastroenterol Motil. 2012;24(6):521–30, e248.

15. Matto J, Maunuksela L, Kajander K, Palva A, Korpela R, Kassinen A, et al. Composition and temporal stability of gastrointestinal microbiota in irritable bowel syndrome—a longitudinal study in IBS and control subjects. FEMS Immunol Med Microbiol. 2005;43(2):213–22.

16. Rea K, O’Mahony SM, Dinan TG, Cryan JF. The Role of the Gastrointestinal Microbiota in Visceral Pain. Handb Exp Pharmacol. 2017;239:269–87.

17. Alexeev EE, Lanis JM, Kao DJ, Campbell EL, Kelly CJ, Battista KD, et al. Microbiota-Derived Indole Metabolites Promote Human and Murine Intestinal Homeostasis through Regulation of Interleukin-10 Receptor. Am J Pathol. 2018;188(5):1183–94.

18. Lamas B, Richard ML, Leducq V, Pham HP, Michel ML, Da Costa G, et al. CARD9 impacts colitis by altering gut microbiota metabolism of tryptophan into aryl hydrocarbon receptor ligands. Nat Med. 2016;22(6):598–605.

19. Neavin DR, Liu D, Ray B, Weinshilboum RM. The Role of the Aryl Hydrocarbon Receptor (AHR) in Immune and Inflammatory Diseases. Int J Mol Sci. 2018;19(12).

20. Postal BG, Ghezzal S, Aguanno D, Andre S, Garbin K, Genser L, et al. AhR activation defends gut barrier integrity against damage occurring in obesity. Mol Metab. 2020;39:101007.

21. Hockley JRF, Taylor TS, Callejo G, Wilbrey AL, Gutteridge A, Bach K, et al. Single-cell RNAseq reveals seven classes of colonic sensory neuron. Gut. 2019;68(4):633–44.

22. Sheu ML, Pan LY, Sheehan J, Yang MY, Pan HC. Modulation of Aryl Hydrocarbon Receptor Expression Alleviated Neuropathic Pain in a Chronic Constriction Nerve Injury Animal Model. Int J Mol Sci. 2022;23(19).

23. Lapointe TK, Basso L, Iftinca MC, Flynn R, Chapman K, Dietrich G, et al. TRPV1 sensitization mediates postinflammatory visceral pain following acute colitis. Am J Physiol Gastrointest Liver Physiol. 2015;309(2):G87–99.

24. Park YH, Kim N, Shim YK, Choi YJ, Nam RH, Choi YJ, et al. Adequate Dextran Sodium Sulfate-induced Colitis Model in Mice and Effective Outcome Measurement Method. J Cancer Prev. 2015;20(4):260–7.

25. Erben U, Loddenkemper C, Doerfel K, Spieckermann S, Haller D, Heimesaat MM, et al. A guide to histomorphological evaluation of intestinal inflammation in mouse models. Int J Clin Exp Pathol. 2014;7(8):4557–76.

26. Vicentini FA, Mathews AJ, Pittman QJ, Swain MG, Sharkey KA, Hirota SA. Behavioural adaptations after antibiotic treatment in male mice are reversed by activation of the aryl hydrocarbon receptor. Brain Behav Immun. 2021;98:317–29.

27. Rondeau LE, Da Luz BB, Haas D, Muppidi P, Wang X, Dang R, et al. Dietary Tryptophan Enhances Aryl Hydrocarbon Receptor Activation and Reduces Colitis Through Microbial Metabolism. Cell Mol Gastroenterol Hepatol. 2026;20(9):101811.

28. Gu Z, Pei W, Shen Y, Wang L, Zhu J, Zhang Y, et al. Akkermansia muciniphila and its outer protein Amuc_1100 regulates tryptophan metabolism in colitis. Food Funct. 2021;12(20):10184–95.

29. Aoki R, Aoki-Yoshida A, Suzuki C, Takayama Y. Indole-3-Pyruvic Acid, an Aryl Hydrocarbon Receptor Activator, Suppresses Experimental Colitis in Mice. J Immunol. 2018;201(12):3683–93.

30. Scott SA, Fu J, Chang PV. Dopamine receptor D2 confers colonization resistance via gut microbial metabolites. bioRxiv. 2023.

31. Chen C, Cao Z, Lei H, Zhang C, Wu M, Huang S, et al. Microbial Tryptophan Metabolites Ameliorate Ovariectomy-Induced Bone Loss by Repairing Intestinal AhR-Mediated Gut-Bone Signaling Pathway. Adv Sci (Weinh). 2024;11(36):e2404545.

32. Tramullas M, Collins JM, Fitzgerald P, Dinan TG, SM OM, Cryan JF. Estrous cycle and ovariectomy-induced changes in visceral pain are microbiota-dependent. iScience. 2021;24(8):102850.

33. Hu W, Sorrentino C, Denison MS, Kolaja K, Fielden MR. Induction of cyp1a1 is a nonspecific biomarker of aryl hydrocarbon receptor activation: results of large scale screening of pharmaceuticals and toxicants in vivo and in vitro. Mol Pharmacol. 2007;71(6):1475–86.

34. Nieves KM, Hirota SA, Flannigan KL. Xenobiotic receptors and the regulation of intestinal homeostasis: harnessing the chemical output of the intestinal microbiota. Am J Physiol Gastrointest Liver Physiol. 2022;322(2):G268–G81.

35. Obata Y, Castano A, Boeing S, Bon-Frauches AC, Fung C, Fallesen T, et al. Neuronal programming by microbiota regulates intestinal physiology. Nature. 2020;578(7794):284–9.

36. Murata Y, Yasaka T, Takano M, Ishihara K. Neuronal and glial expression of inward rectifier potassium channel subunits Kir2.x in rat dorsal root ganglion and spinal cord. Neurosci Lett. 2016;617:59–65.

37. Lonnfors S, Vermeire S, Greco M, Hommes D, Bell C, Avedano L. IBD and health-related quality of life --discovering the true impact. J Crohns Colitis. 2014;8(10):1281–6.

38. Siegel CA, MacDermott RP. Is chronic pain an extraintestinal manifestation of IBD? Inflamm Bowel Dis. 2009;15(5):769–71.

39. Minderhoud IM, Oldenburg B, Wismeijer JA, van Berge Henegouwen GP, Smout AJ. IBS-like symptoms in patients with inflammatory bowel disease in remission; relationships with quality of life and coping behavior. Dig Dis Sci. 2004;49(3):469–74.

40. Willing BP, Dicksved J, Halfvarson J, Andersson AF, Lucio M, Zheng Z, et al. A pyrosequencing study in twins shows that gastrointestinal microbial profiles vary with inflammatory bowel disease phenotypes. Gastroenterology. 2010;139(6):1844–54 e1.

41. Martinez C, Antolin M, Santos J, Torrejon A, Casellas F, Borruel N, et al. Unstable composition of the fecal microbiota in ulcerative colitis during clinical remission. Am J Gastroenterol. 2008;103(3):643–8.

42. Morgan XC, Tickle TL, Sokol H, Gevers D, Devaney KL, Ward DV, et al. Dysfunction of the intestinal microbiome in inflammatory bowel disease and treatment. Genome Biol. 2012;13(9):R79.

43. Scott SA, Fu J, Chang PV. Microbial tryptophan metabolites regulate gut barrier function via the aryl hydrocarbon receptor. Proc Natl Acad Sci U S A. 2020;117(32):19376–87.

44. McKillop D, Case DE. Mutagenicity, carcinogenicity and toxicity of beta-naphthoflavone, a potent inducer of P448. Biochem Pharmacol. 1991;41(1):1–7.

45. Mimura J, Fujii-Kuriyama Y. Functional role of AhR in the expression of toxic effects by TCDD. Biochim Biophys Acta. 2003;1619(3):263–8.

46. Metidji A, Omenetti S, Crotta S, Li Y, Nye E, Ross E, et al. The Environmental Sensor AHR Protects from Inflammatory Damage by Maintaining Intestinal Stem Cell Homeostasis and Barrier Integrity. Immunity. 2018;49(2):353–62 e5.

47. Lamas B, Natividad JM, Sokol H. Aryl hydrocarbon receptor and intestinal immunity. Mucosal Immunol. 2018;11(4):1024–38.

48. Gangadharan V, Kuner R. Pain hypersensitivity mechanisms at a glance. Dis Model Mech. 2013;6(4):889–95.

49. Hucho T, Levine JD. Signaling pathways in sensitization: toward a nociceptor cell biology. Neuron. 2007;55(3):365–76.

50. Basbaum AI, Bautista DM, Scherrer G, Julius D. Cellular and molecular mechanisms of pain. Cell. 2009;139(2):267–84.

51. Brierley SM, Hibberd TJ, Spencer NJ. Spinal Afferent Innervation of the Colon and Rectum. Front Cell Neurosci. 2018;12:467.

52. Defaye M, Abdullah NS, Iftinca M, Hassan A, Agosti F, Zhang Z, et al. Gut-innervating TRPV1+ Neurons Drive Chronic Visceral Pain via Microglial P2Y12 Receptor. Cell Mol Gastroenterol Hepatol. 2022;13(4):977–99.

53. Meynier M, Baudu E, Rolhion N, Defaye M, Straube M, Daugey V, et al. AhR/IL-22 pathway as new target for the treatment of post-infectious irritable bowel syndrome symptoms. Gut Microbes. 2022;14(1):2022997.

54. Wang X, Hu M, Wu W, Lou X, Gao R, Ma T, et al. Indole derivatives ameliorated the methamphetamine-induced depression and anxiety via aryl hydrocarbon receptor along “microbiota-brain” axis. Gut Microbes. 2025;17(1):2470386.

