## Supplemental Tables 1 and 2 for "ACTIVATION OF THE ARYL HYDROCARBON RECEPTOR REDUCES VISCERAL HYPERSENSITIVITY IN A MOUSE MODEL OF COLITIS IN REMISSION"

### Supplemental table 1

♂

| 45 mmHg | Control Veh+Veh | Control GNF+Veh | Control Veh+IPyA | Control GNF+IPyA | Post-colitis Veh+Veh | Post-colitis GNF+Veh | Post-colitis Veh+IPyA | Post-colitis GNF+IPyA |
|---|---|---|---|---|---|---|---|---|
| Control Veh+Veh | | | | | | | | |
| Control GNF+Veh | 0.0414 | | | | | | | |
| Control Veh+IPyA | ns | 0.0093 | | | | | | |
| Control GNF+IPyA | ns | ns | ns | | | | | |
| Post-colitis Veh+Veh | <0.0001 | ns | 0.0242 | 0.0072 | | | | |
| Post-colitis GNF+Veh | 0.0178 | ns | ns | ns | ns | | | |
| Post-colitis Veh+IPyA | ns | ns | ns | ns | <0.0001 | 0.0035 | | |
| Post-colitis GNF+IPyA | ns | ns | ns | ns | 0.0006 | ns | ns | |

### Supplemental table 2

♂

| 60 mmHg | Control Veh+Veh | Control GNF+Veh | Control Veh+IPyA | Control GNF+IPyA | Post-colitis Veh+Veh | Post-colitis GNF+Veh | Post-colitis Veh+IPyA | Post-colitis GNF+IPyA |
|---|---|---|---|---|---|---|---|---|
| Control Veh+Veh | | | | | | | | |
| Control GNF+Veh | ns | | | | | | | |
| Control Veh+IPyA | ns | ns | | | | | | |
| Control GNF+IPyA | 0.0432 | ns | ns | | | | | |
| Post-colitis Veh+Veh | <0.0001 | ns | 0.0011 | ns | | | | |
| Post-colitis GNF+Veh | 0.0290 | ns | ns | ns | ns | | | |
| Post-colitis Veh+IPyA | ns | 0.0498 | ns | 0.0083 | <0.0001 | 0.0052 | | |
| Post-colitis GNF+IPyA | ns | ns | ns | ns | ns | ns | 0.0408 | |
