## Supplemental Tables 3 and 4 for "ACTIVATION OF THE ARYL HYDROCARBON RECEPTOR REDUCES VISCERAL HYPERSENSITIVITY IN A MOUSE MODEL OF COLITIS IN REMISSION"

### Supplemental table 3

♀

| 45 mmHg | Control Veh+Veh | Control GNF+Veh | Control Veh+IPyA | Control GNF+IPyA | Post-colitis Veh+Veh | Post-colitis GNF+Veh | Post-colitis Veh+IPyA | Post-colitis GNF+IPyA |
|---|---|---|---|---|---|---|---|---|
| Control Veh+Veh | | | | | | | | |
| Control GNF+Veh | ns | | | | | | | |
| Control Veh+IPyA | ns | ns | | | | | | |
| Control GNF+IPyA | ns | ns | ns | | | | | |
| Post-colitis Veh+Veh | ns | ns | ns | ns | | | | |
| Post-colitis GNF+Veh | ns | ns | ns | ns | ns | | | |
| Post-colitis Veh+IPyA | ns | ns | ns | 0.0352 | 0.0143 | 0.0075 | | |
| Post-colitis GNF+IPyA | ns | ns | ns | ns | ns | ns | 0.0195 | |

### Supplemental table 4

♀

| 60 mmHg | Control Veh+Veh | Control GNF+Veh | Control Veh+IPyA | Control GNF+IPyA | Post-colitis Veh+Veh | Post-colitis GNF+Veh | Post-colitis Veh+IPyA | Post-colitis GNF+IPyA |
|---|---|---|---|---|---|---|---|---|
| Control Veh+Veh | | | | | | | | |
| Control GNF+Veh | 0.0286 | | | | | | | |
| Control Veh+IPyA | ns | ns | | | | | | |
| Control GNF+IPyA | 0.0032 | ns | ns | | | | | |
| Post-colitis Veh+Veh | 0.0025 | ns | ns | ns | | | | |
| Post-colitis GNF+Veh | ns | ns | ns | ns | ns | | | |
| Post-colitis Veh+IPyA | ns | ns | ns | 0.0130 | 0.0115 | ns | | |
| Post-colitis GNF+IPyA | 0.0446 | ns | ns | ns | ns | ns | ns | |
