## Supplementary figures and images for "ACTIVATION OF THE ARYL HYDROCARBON RECEPTOR REDUCES VISCERAL HYPERSENSITIVITY IN A MOUSE MODEL OF COLITIS IN REMISSION"

### Supplemental Figure 1

Supplemental Figure 1

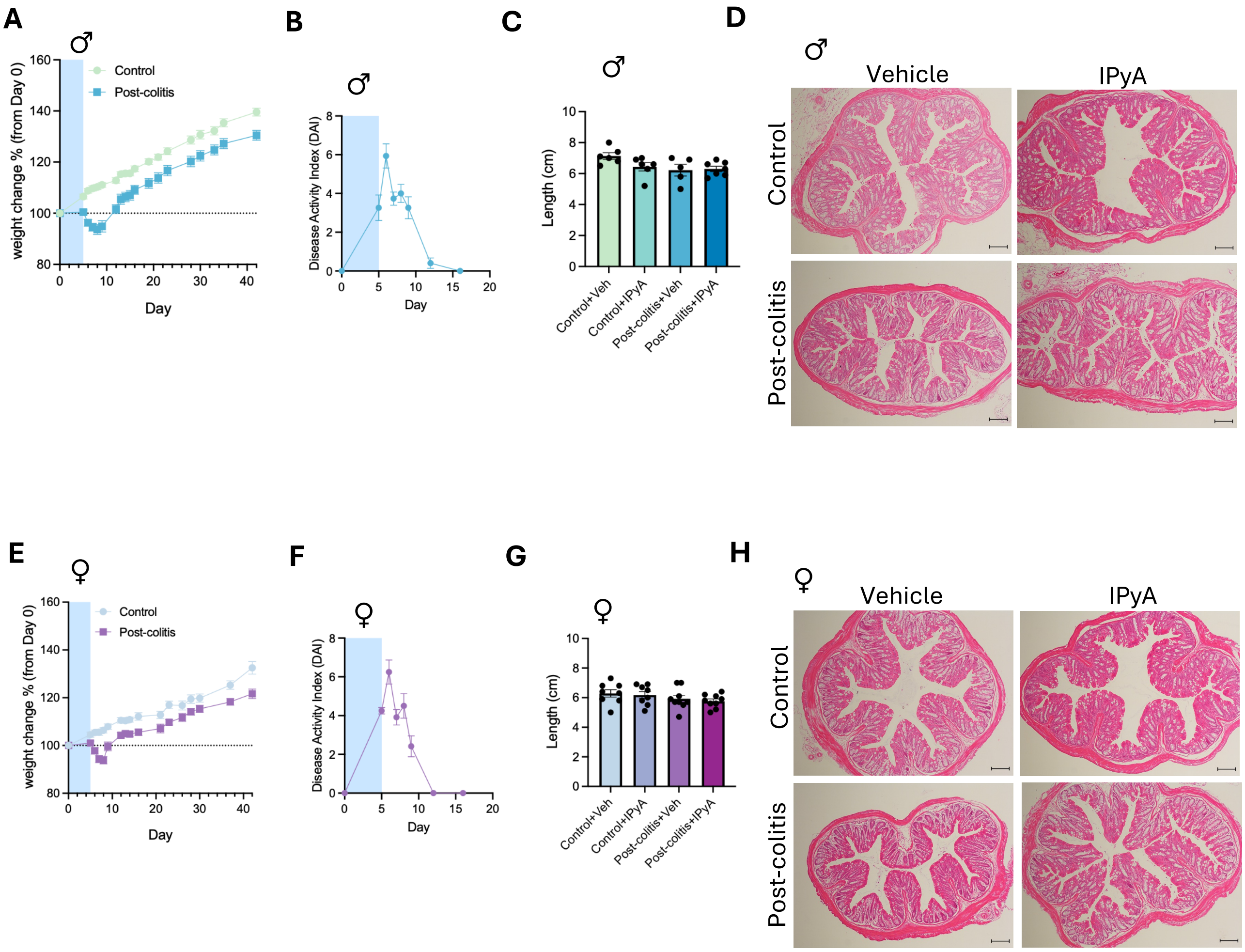

### Supplemental Figure 2

#### Supplemental Figure 2

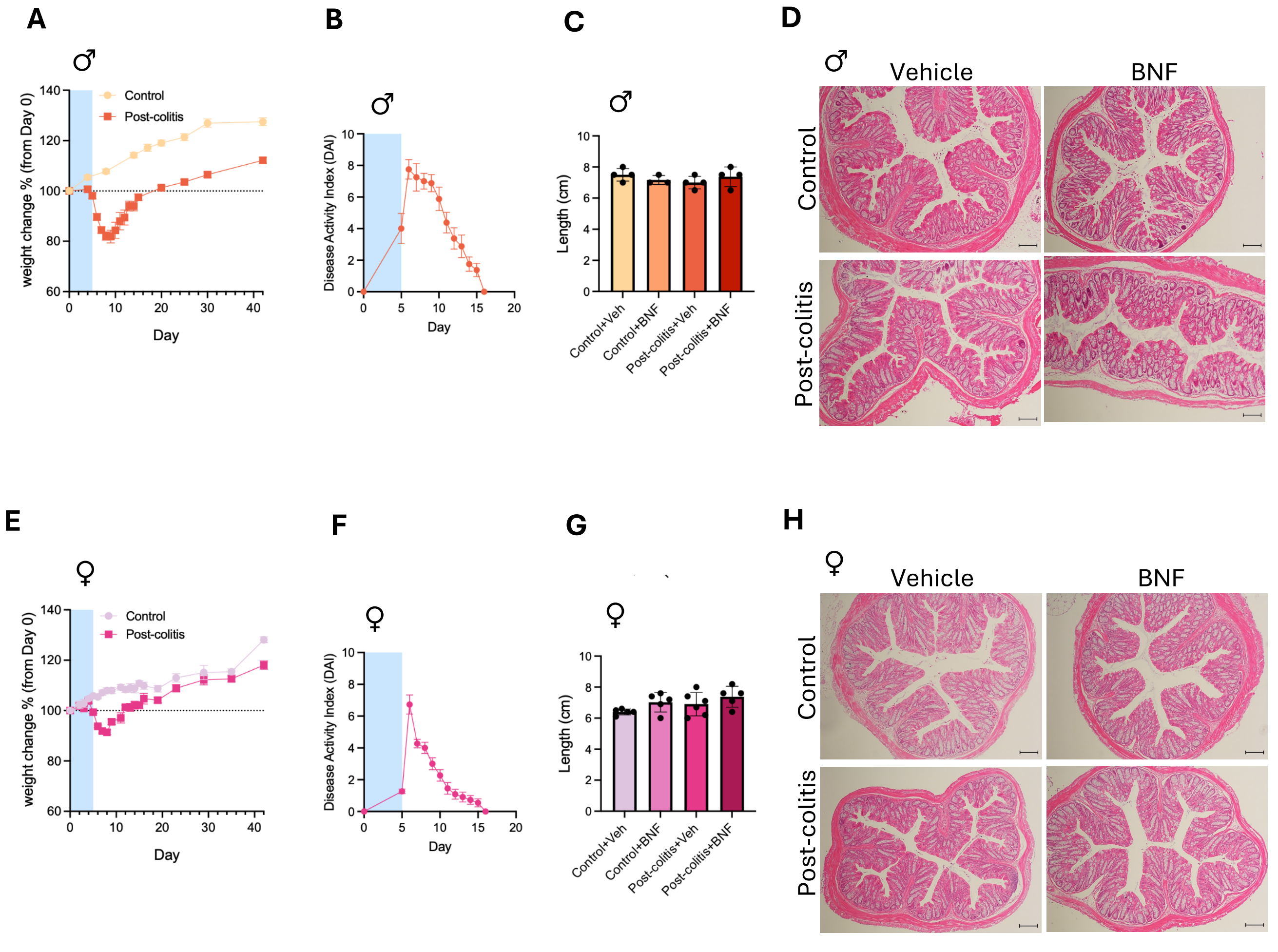

### Supplemental Figure 3

#### Supplemental Figure 3

**A**

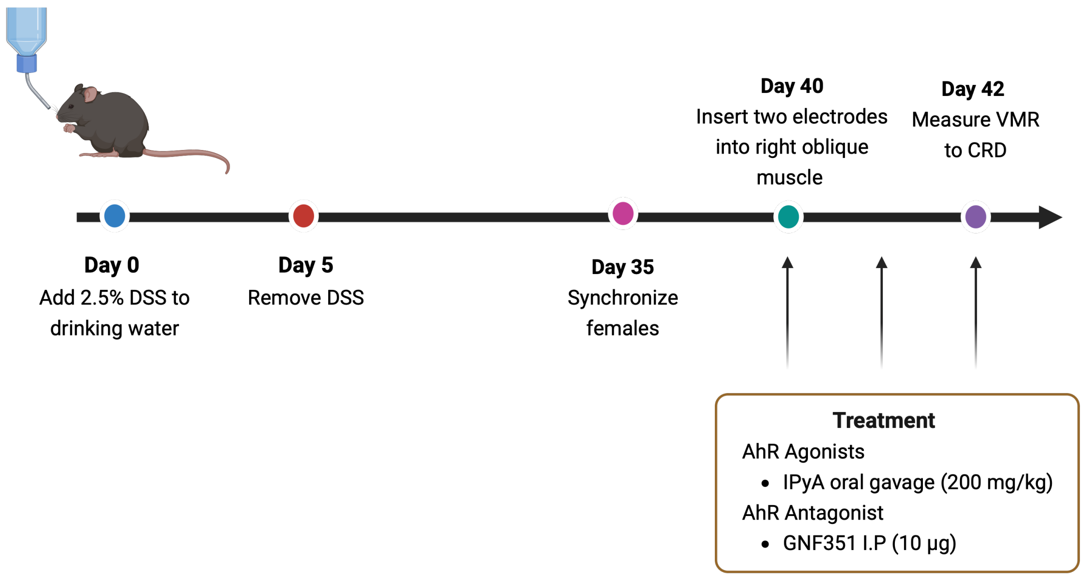


**B**

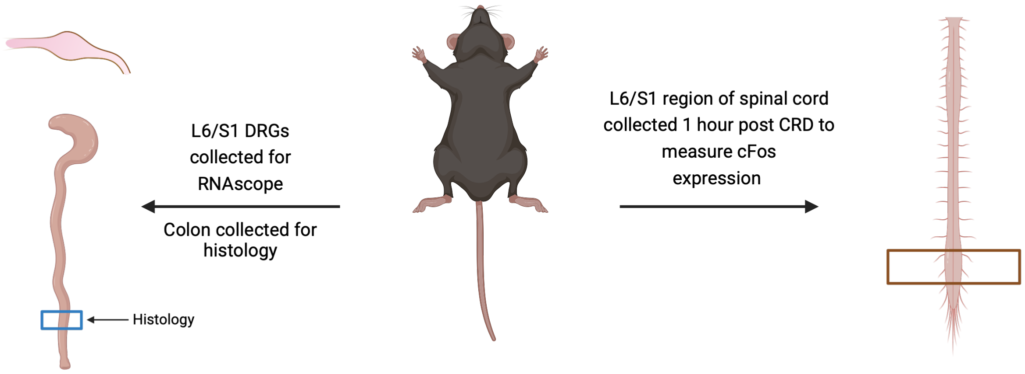

### Supplemental Figure 4

#### Supplemental Figure 4

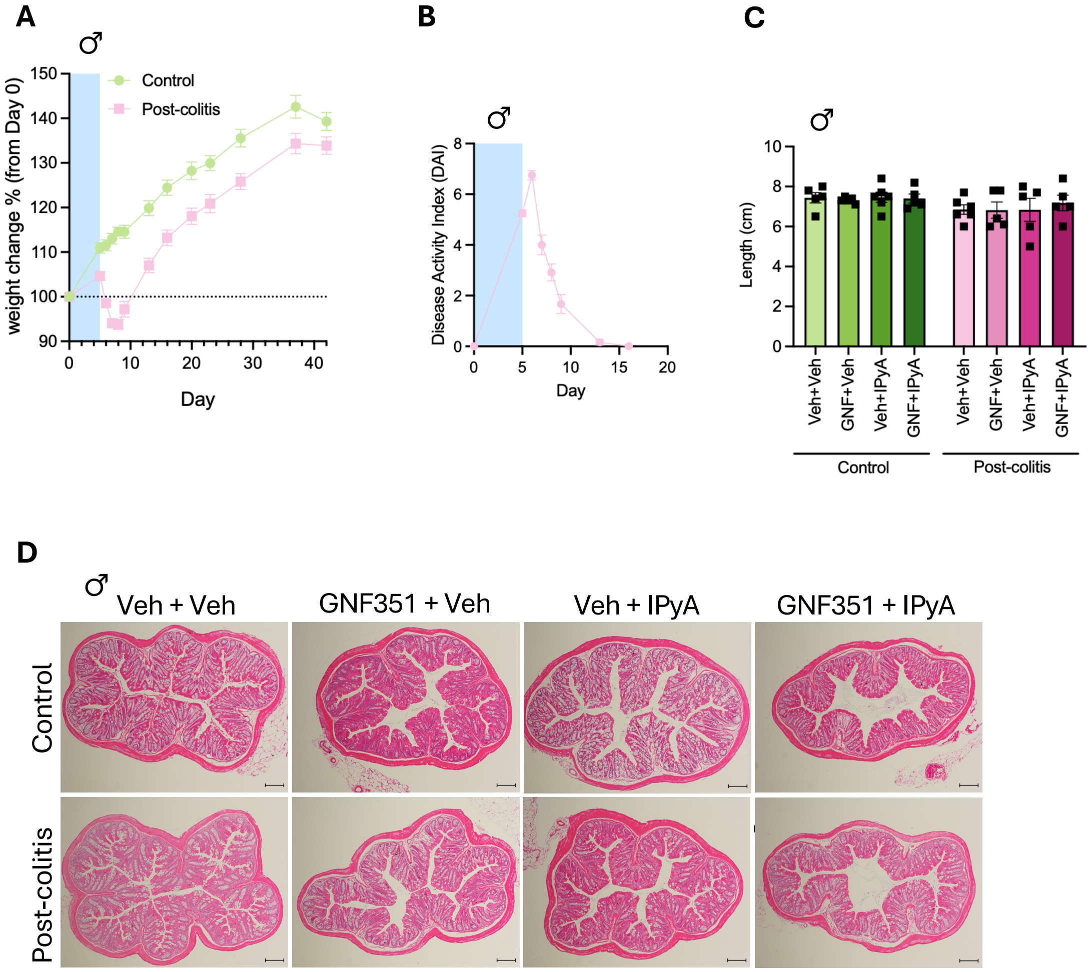

### Supplemental Figure 5

Supplemental Figure 5

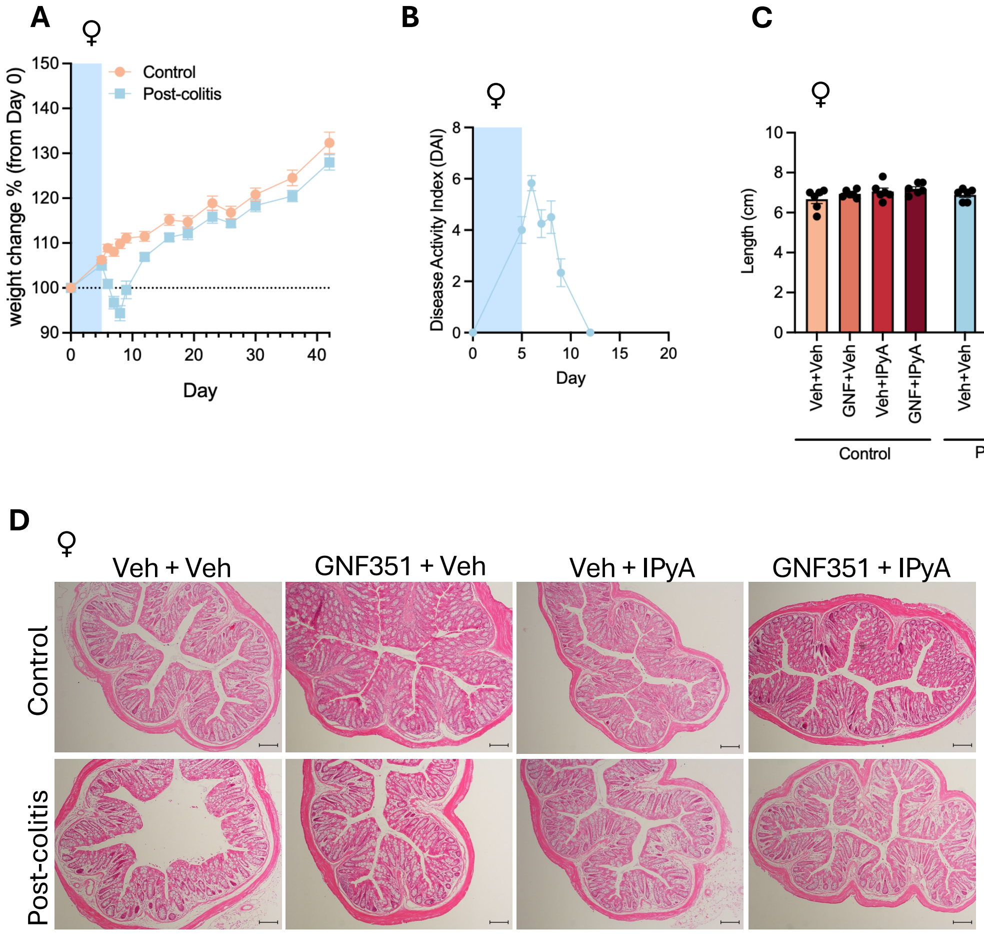

### Supplemental Figure 6

#### Supplemental Figure 6

**A**

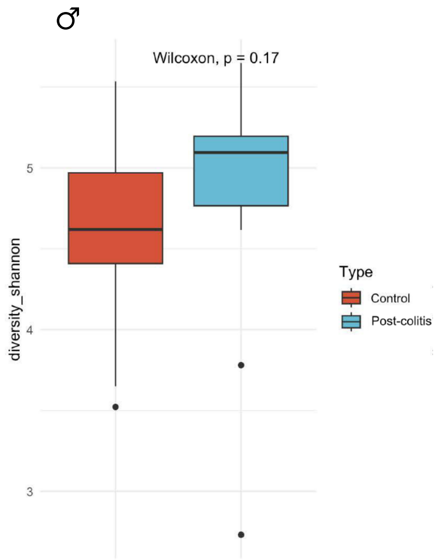


**B**

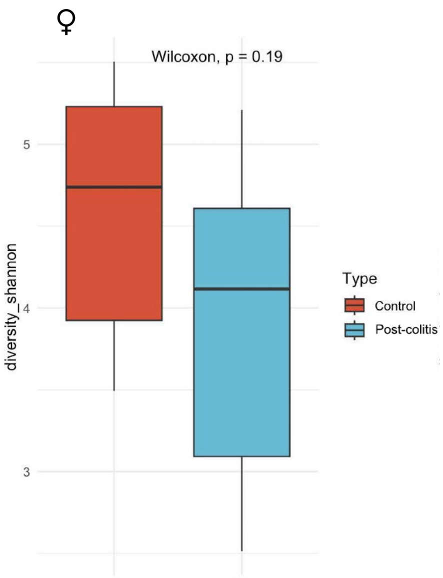

### Supplemental Figure 7

#### Supplemental Figure 7

**A**

♂

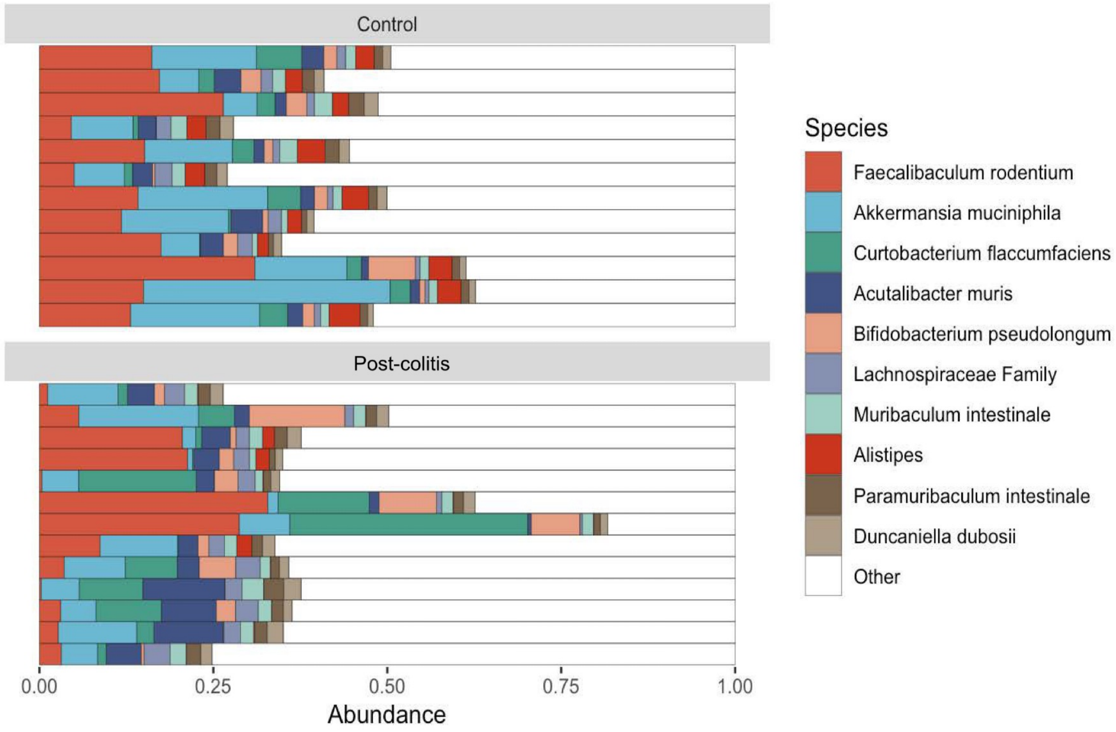


**B**

♀

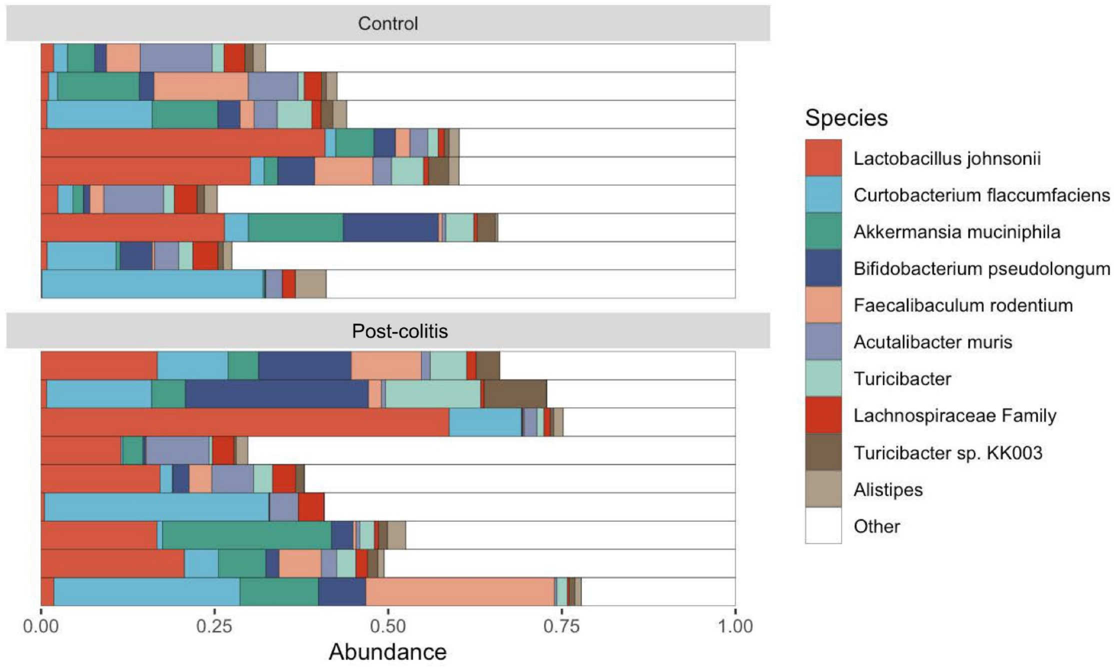
